# Brain-derived neurotrophic factor (BDNF), estrogen receptors, and mossy cells

**DOI:** 10.64898/2026.10.16.759700

**Authors:** Chiara Criscuolo, Monarchsinh Rathod, Fang Jie, Helen E. Scharfman

## Abstract

Dentate gyrus (DG) function depends on the neurotrophin brain-derived neurotrophic factor (BDNF) which shows robust synthesis in the principal cells, the granule cells (GCs). However, BDNF protein is present in the inner molecular layer (IML), where most afferents to GCs are from hilar mossy cells (MCs). Therefore, we asked whether MCs express BDNF mRNA and protein. Using adult mice, we found that MC BDNF mRNA and protein were expressed in the majority of MC somata and few other hilar neurons. Interestingly, IML BDNF was significantly greater in the ventral compared to dorsal DG. Additionally, dorsal IML BDNF was significantly higher in females on proestrous morning, when serum levels of estrogen peak, relative to metestrous morning, when estrogen is lower. Consistent with this observation, ventral MC somata, which project to the dorsal IML, showed more BDNF mRNA on proestrous morning than metestrous morning. Next, we asked if strong IML BDNF protein expression on proestrous morning was due to estrogen action at estrogen receptors (ERs) because estrogen binding to ERs can increase BDNF synthesis. We found that MCs have high expression of all ERs: ERα, ERβ and GPER1 in both sexes. When sexes were pooled, the percent of MCs labeled by ERβ and GPER1 exceeded the percent of MCs expressing ERα. When proestrous females were examined, GPER1-labeled hilar cells were greater than hilar cells labeled by ERα. The results demonstrate robust MC somatic expression of BDNF, relatively high ventral IML expression of BDNF, and estrous cycle dependence. The results also highlight robust MC ER expression, especially ERβ and GPER1, making the hilus a previously underestimated site of estrogen and BDNF action.

**HIGHLIGHTS:**

- BDNF mRNA and protein are expressed in most MCs and their axons in both sexes.
- Ventral MC axons in the distal inner molecular layer (IML) exhibit more BDNF protein than dorsal MC axons.
- Ventral MCs show relatively strong BDNF mRNA on proestrous morning compared to metestrous females.
- MCs express all 3 estrogen receptors (ERα, ERβ, GPER1) in both sexes, allowing estrogen to influence BDNF by MC ERs.

**GRAPHICAL ABSTRACT:** <u>Top:</u> A circuit diagram of the hippocampus shows cell layers in grey and the principal cell types (dentate gyrus granule cells, CA3 pyramidal cells, CA1 pyramidal cells). In the past. BDNF protein was considered to be produced mainly in granule cells and transported to their axons.

<u>Bottom:</u> The implications of the present study are illustrated schematically. For example, hilar mossy cells (MCs) express BDNF mRNA and protein, and express all estrogen receptors (ERs): ERα, ERβ, GPER1. Therefore, MC ERs have the potential to play important roles in BDNF and estrogen action in the dentate gyrus.

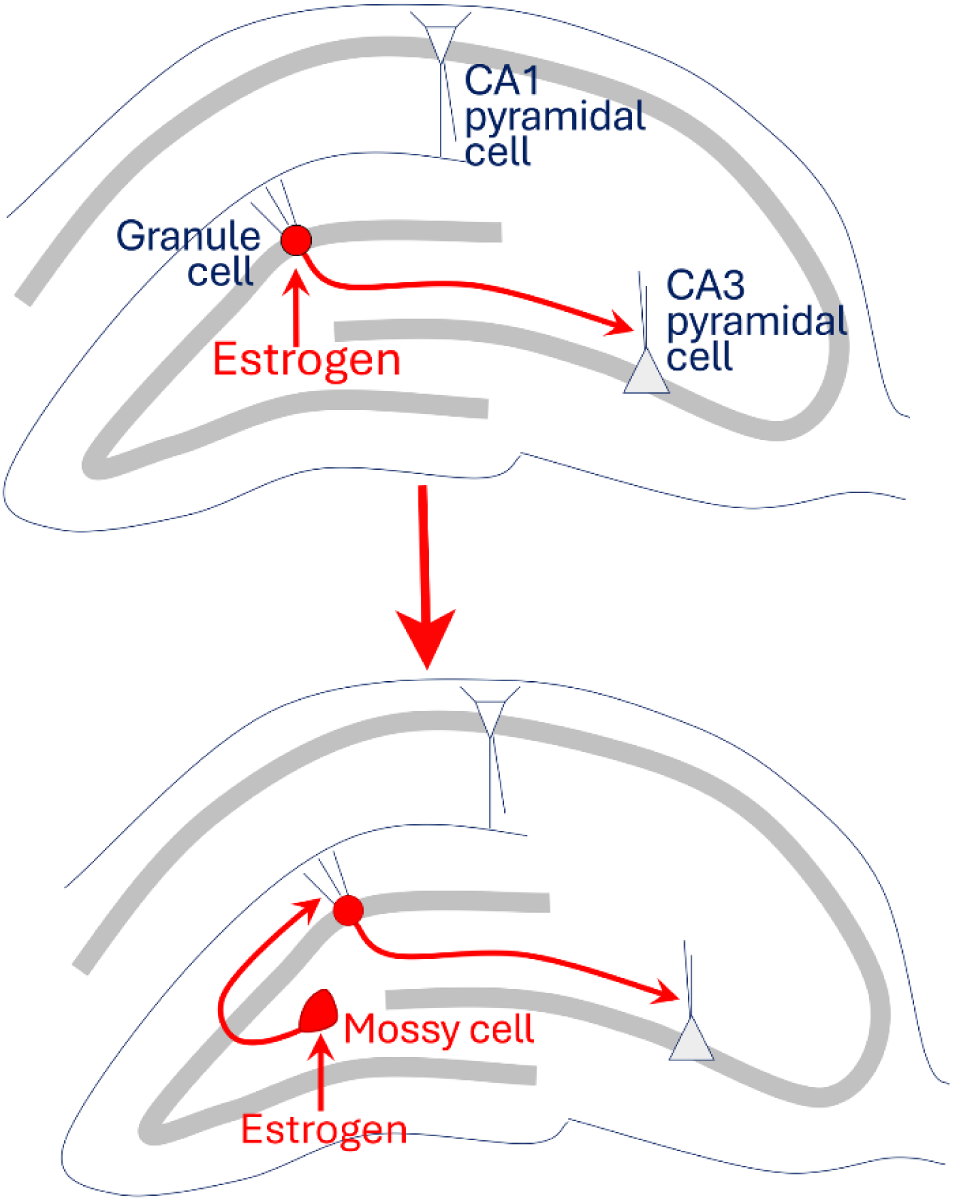

## INTRODUCTION

Brain-derived neurotrophic factor (BDNF) is a member of the neurotrophin family and binds primarily to tropomyosin receptor kinase B (TrkB) and, with lower affinity, p75 neurotrophin receptors (p75NTR) [1,2]. In the hippocampal dentate gyrus (DG), BDNF has many functions related to structure and function of the principal cells, glutamatergic granule cells (GCs). Without BDNF, adverse effects occur on GC dendritic spines, dendrites, and axons [3,4]. Perhaps for this reason, functions considered to critically depend on GCs, such as the regulation of mood, are impaired after selective deletion of BDNF in the DG [5]. BDNF is also critical to the normal maturation of GCs that are born in adulthood (adult-born GCs), which regulate mood as well as DG-dependent cognitive functions [6–14].

For these reasons, most attention has focused on the role of BDNF in GCs rather than other DG cell types. It is assumed that BDNF is derived from the GCs themselves, since BDNF mRNA and protein are expressed at high levels in GCs [15,16]. Interestingly, most BDNF is anterogradely transported to the GC axons, the mossy fibers (MFs; [15–17]). Anterograde transport is so robust that an antibody to BDNF mainly detects BDNF protein in the MFs rather than GC somata [15–17]. BDNF is stored in dense core vesicles in the MF terminals [17] and is released preferentially upon high frequency stimulation [18], leading to long-term potentiation (LTP) of MF transmission to CA3 neurons [19–21]. Exposure of hippocampal slices to recombinant BDNF also leads to potentiation of the MF pathway without LTP of other inputs to CA3 [22], suggesting a unique role of BDNF to potentiate the MF input. The effects of BDNF on MF transmission are significant for hippocampal-dependent memory, where the MF pathway plays a critical role [23,24]. BDNF also mediates a form of long-term depression (LTD) of MF transmission to CA3 neurons [25], which is surprising given that BDNF induces MF LTP. One explanation is that MF-LTP is dependent on TrkB [19] whereas BDNF-dependent MF LTD is not [25]. It is also potentially important that exposure to recombinant BDNF in vivo potentiates the major cortical input to GCs, the perforant path (PP) input from entorhinal cortex [26]. BDNF also has been shown to regulate GABAergic inhibition of GCs [27], as well as astrocytes and microglia [28].

In contrast to these studies, less is known about BDNF protein expression in mossy cells (MCs) of the DG. MCs are the major cell type in the hilus, are the only glutamatergic DG cell type besides GCs, and project primarily to the proximal dendrites of GCs in the inner molecular layer (IML; [29–31]). The projection mainly terminates distant to the MC soma, both ipsi- and contralaterally [29,32]. However, MCs also innervate DG GABAergic neurons [29,32]. MC BDNF is potentially important because its release could induce synaptic plasticity of MC-GC or MC-GABAergic neuron synapses. It is important to understand more about MC BDNF in light of recent studies showing that BDNF release from GCs was required for a form of LTP at the MC-GC synapse [33–35]. It was shown that BDNF protein in dendrites of GCs was released, traveled retrogradely to MC terminals, and acted on adenosine receptors on MC terminals to regulate MC glutamate release.

From past studies of BDNF distribution in the brain, it was shown that BDNF protein is expressed in the IML [15,16]. Therefore, MCs appear to express BDNF in their axons in the IML. However, somatic expression of BDNF protein in MCs has not been shown, probably because BDNF protein in hilar MFs of GCs is so high that it obscures the detection of BDNF in MCs. From studies of gene expression by transcriptomics, it has been shown that MCs have BDNF mRNA [36], although the extent of BDNF synthesis and its variability throughout the MC population has not been studied. This leads to several questions. First, to what extent do MC somata express BDNF mRNA and protein? Second, does MC BDNF vary along the septotemporal axis? The septotemporal axis is important because functions of dorsal and ventral DG differ [37–41].

Another question is whether MC BDNF varies with sex and the estrous cycle. Sex differences might be expected because GC BDNF protein is greater in female rats compared to males [42]. Fluctuations in estrogen could influence BDNF because MF BDNF in rats is greatest after serum levels of 17β-estradiol peak on proestrous morning of the estrous cycle [42–44]. It has been suggested that 17β-estradiol increases BDNF by binding to estrogen receptors (ERs) that translocate to the nucleus and activate an estrogen-sensitive response element (ERE) on the BDNF gene, or ERs act at the BDNF promoter, increasing BDNF synthesis [45,46]. Alternatively, in cortical circuits, 17β-estradiol may promote principal neuron activity by actions on GABAergic neurons [47,48], leading to increased BDNF in principal neurons by activity-dependent BDNF transcription [49].

Interestingly, it is not yet clear that MCs express ERs. Initial studies of ERα mRNA showed high levels in hilar neurons [50], but a subsequent publication suggested that few hilar neurons express ERα [51]. Another study of ERα showed hilar neurons expressed ERα, but most were likely to be GABAergic based on their ultrastructure and location [52]. Regarding ERβ, initially it did not appear to be robust in hilar neurons [50]. However, later it was shown that hilar ERβ protein was present [53,54]. GPER1 protein also appears to be robust in hilar neurons [55]. A diagram of the DG circuit showed MCs had ERα and ERβ [51], but to our knowledge, double labeling with MC markers has not been performed. Robust ERs on MCs might be functionally important because estrogen promotes lesion-induced sprouting of axons into the IML [56], which may arise from MCs. Estrogen actions on MCs may also be important to protect them from toxicity, because MCs are highly vulnerable to seizures, ischemia, and brain injury [57]. Indeed, past studies have shown that estrogen protects hilar neurons [58–60], although mechanisms were suggested to be glial [60] rather than due to ERs on MCs. MC ERs may also contribute to social behaviors because estrogen plays an important role in regulating these behaviors [61]. Indeed, MCs appear to be essential for social behavior, although the studies to date focused on MC oxytocin receptors [62] rather than ERs and BDNF.

For these reasons, we examined BDNF mRNA and protein expression in MC somata and their axons in the IML in male and female adult mice. Next, the septotemporal axis was examined. Comparisons were made between males and two groups of females: those perfused on metestrous morning, the time of the estrous cycle when serum levels of 17β-estradiol are relatively low, and females on proestrous morning, a time when serum levels of 17β-estradiol peak. Proestrous females had higher ventral MC BDNF mRNA and greater dorsal IML BDNF than metestrous females, suggesting estrogen might increase BDNF synthesis, as it does in GCs [42,63]. Therefore, we determined if MCs express ERs and found robust expression of all 3 ERs: ERα, ERβ and G-protein coupled ER 1 (GPER1). Together the results demonstrate sex and estrous-cycle dependent BDNF expression in MCs and its variation along the septotemporal axis. The results also show that there is robust ER and BDNF expression not only in GCs but also in MCs, suggesting that there is much more to BDNF action and its regulation in the DG than just the GCs.

## MATERIALS AND METHODS

The experimental procedures were performed according to National Institutes of Health guidelines and approved by the Institutional Animal Care and Use Committee at the Nathan Kline Institute.

### I. Terminology

*Dorsal* refers to the part of the hippocampus closest to the septum and in the dorsal half of the forebrain. *Ventral* refers to the part of the hippocampus closest to the temporal pole and is in the ventral half of the forebrain. This distinction is worth noting because some investigators refer to the caudal, dorsal part of hippocampus as ventral.

Throughout the manuscript, the word *“estrogen”* refers to all 3 forms (17β-estradiol, estrone, estriol) analogous to the term “*estrogens*” used elsewhere [64]. When 17β-estradiol was used specifically, the term *17β-estradiol* is used instead of *estrogen*.

### II. Experimental design

Adult female and male mice were compared to assess BDNF mRNA and protein in MCs. Vaginal cytology was used to distinguish females at 2 phases of their estrous cycle, one with relatively high levels of serum 17β-estradiol (proestrous morning), and the other with relatively low levels (metestrous morning; for details see below). Thus, three groups were compared: males, females with relatively high levels of estradiol, and females with relatively low levels. After perfusion-fixation, the hippocampus was sectioned in a way that allowed visualization of the DG throughout the septotemporal axis: one hemisphere was sectioned coronally and the opposite hemisphere was sectioned horizontally. Coronal sections were used to assess the dorsal part of the hippocampus and horizontal sections were used for the ventral DG. This approach was employed because the ventral molecular layer and hilus of the DG are much more distinct in the horizontal plane compared to coronal sections. Because 17β-estradiol can increase BDNF protein synthesis by actions at ERs [45,46], we also asked if MCs expressed ERα, ERβ, or GPER1 protein. Then we asked if BDNF protein was increased by neuronal activity because 17β-estradiol can increase neuronal activity [65–67], and some BDNF transcripts are activity-dependent [49,68]. Therefore, we double labeled sections with antibodies to a MC marker, calretinin, and a marker of neuronal activity, c-Fos.

### III. Animal care

Mice were housed in standard mouse cages with corn cob bedding and a 12-hour light-dark cycle (lights on: 7:00 a.m., lights off, 7:00 p.m.). Mice had food and water *ad libitum*. During gestation and until weaning, mice were fed chow formulated for breeding (Formulab diet 5008; LabDiet). After weaning they were fed a standard rodent chow (Rodent diet 5001, LabDiet). It should be noted that the 5001 chow contains soy and alfalfa, which are sources of phytoestrogens [69,70]. After weaning, mice were housed with littermates of the same sex (maximum four mice per cage).

#### Breeding and genotyping

Wild-type (WT) mice were selected from a colony of Tg2576 transgenic mice, which express the Swedish mutation in human amyloid precursor protein 695 driven by the hamster prion protein promoter, and simulate features of Alzheimer’s disease [71]. They were bred in-house from male heterozygous Tg2576 and female non-transgenic mice (C57BL6/SJL F1 hybrid, Stock# 100012, Jackson Labs). Only non-transgenic (WT) offspring were used in this study. Mice were weaned at 23-25 days of age. Genotype was determined using an assay for APP695 by the Mouse Genotyping core facility at New York University Grossman School of Medicine. WT mice were identified by the absence of detectable APP695.

#### Determination of estrous cycle phase in female mice

There are sequential phases of the rodent estrous cycle: proestrus, estrus, and 2 stages of diestrus. Diestrus has an initial 24 hrs called diestrus 1 (also called metestrus) and a subsequent 24 hrs called diestrus 2. The estrous cycle phase with the highest serum level of 17β-estradiol is mid-morning of proestrus, estimated as 9:00 a.m. in the Sprague-Dawley rat when lights are turned on at 7:00 a.m. [72]. Therefore, mice were euthanized shortly thereafter (at 10:00 a.m.-12:00 p.m.) to capture effects of 17β-estradiol, since the effects would presumably occur shortly after the peak was reached. The phase with the lowest serum levels of 17β-estradiol is metestrus; levels are also low early on diestrus 2 but 17β-estradiol slowly rises later that day [72]. To keep the time of day constant, mice were always perfused between 10:00 a.m. and 12:00 p.m. To be confident of the estrous cycle phase, a sample of the vaginal epithelia was examined microscopically every morning between 9:00 and 11:00 a.m. Monday-Friday for at least 2 weeks prior to experiments. Two weeks was necessary to confirm females had regular estrous cycles. Before starting female estrous cycle monitoring, mice were acclimated to the experimenter through daily handling (1 minute per mouse) for one week. Male mice were handled daily also and the handling of males continued for the two weeks when females were tested for their estrous cycle. To minimize acyclicity, female cages were placed adjacent to cages with males [73].

Characterization of vaginal cells was based on Scharfman et al. [74]. The cell types were defined as follows: leukocytes (round, small cells), epithelial cells (oval, intermediate-size, nucleated cells), and cornified epithelial cells (multipolar, large, nucleated cells). The pattern of vaginal cytology was consistent with a cyclic pattern described previously [75]. This cycle is 4-5 days as follows: one day of proestrus, one day of estrus, and 2-3 days of diestrus. Proestrous morning is characterized by epithelial cells, and some cornified epithelial cells. Estrous morning exhibits a greater percentage of cornified epithelial cells. Diestrous mornings are characterized by a dominance of leukocytes and few epithelial cells. Therefore, we defined proestrous morning as the first day when the vaginal sample was dominated by epithelial cells after 2 or more days with samples that had primarily leukocytes. In contrast, metestrus was defined as the first day that was dominated by leukocytes after 2 or more days with primarily epithelial or cornified epithelial cells. It should be noted that some female mice did not exhibit at least two 4-5 day cyclic patterns prior to experimental use, and they were excluded.

### IV. Anatomy

#### Perfusion-fixation and sectioning

Mice were deeply anesthetized by isoflurane inhalation (NDC#07-893-1389, Patterson Veterinary) followed by intraperitoneal (i.p.) injection of 250 mg/kg urethane (Cat#U2500, Millipore-Sigma). Urethane was made as a 250mg/ml stock solution in 0.9% sodium chloride (NaCl; Cat#S9888; Millipore-Sigma). The abdominal cavity was opened with surgical scissors, followed by the heart cavity. A 26-gauge needle was inserted into the heart, an incision was made in the left atrium, and 10 ml saline (0.9% NaCl in double distilled H_2_O; ddH_2_O) was infused with a peristaltic pump (Minipuls 1; Gilson) followed by 30 ml of cold (4°C) 4% paraformaldehyde (PFA; Cat#19210, Electron Microscopy Sciences) in 0.1 M phosphate buffer (PB; note all buffers were pH 7.4). The brains were removed immediately and postfixed in 4% PFA at 4°C for 3 hours. After postfixation, the brains were washed in 0.1 M PB. Before sectioning, the brains were hemisected. One hemisphere was cut in the coronal plane and the other in the horizontal plane (50 μm-thick sections) using a vibratome (Model# VT1000p, Leica). Similar septotemporal levels were selected and processed together. Sections were collected sequentially in separate wells of a cell culture plate in cryoprotectant solution (30% sucrose, Cat#S8501, Millipore-Sigma, and 30% ethylene glycol, Cat#293237, Millipore-Sigma, diluted in 0.1 M PB). The sections that were not used immediately were stored at −20°C.

#### BDNF immunohistochemistry

BDNF protein was detected with a mouse monoclonal anti-BDNF antibody that has been validated (Mab#9, Developmental Hybridoma Bank [17]). This antibody is similar qualitatively to the one that was used originally in normal rodents that demonstrated robust staining [15,16].

Free-floating sections were first washed in 0.1 M Tris buffered saline (TBS, 3×5 minutes) and then blocked with 3% Mouse-on-Mouse blocking serum (M.O.M; Cat#MKB-2213, Vector Laboratories) in TBS for 1 hour. Sections were incubated for 2 nights at 4°C on a rotator (The Belly Dancer, Stovall) in primary antibody (10 μg/ml) diluted in 3% bovine serum albumin (BSA; Cat#A7906, Millipore-Sigma), 2% donkey serum (DS; D9663, Millipore-Sigma), and 0.3% Triton X-100 (Cat#X-100, Millipore-Sigma) in TBS. Afterwards, sections were incubated in donkey anti–mouse IgG Alexa Fluor 488–conjugated secondary antibody (1:500; Cat#821202, Invitrogen) in 3% DS and 0.3% Triton X-100 in TBS for 1 hour at room temperature (RT), washed in TBS, and then TB. Washes were 3×5 minutes each. Labeled sections were mounted onto gelatin-coated (1% bovine gelatin; Cat#G9391, Millipore-Sigma) glass slides, allowed to dry for a few hours at RT, and coverslipped with fluorescent mounting medium (Citifluor AF1; Cat#17970-25, Electron Microscopy Sciences).

For brightfield microscopy, free-floating sections were first washed in TBS and then treated with 0.25% H_2_O_2_ (Cat#216763, Millipore-Sigma) in TBS for 3 minutes. After washes in TBS, sections were incubated in 0.3% Triton X-100 in TBS for 20 minutes at RT. Next, sections were blocked in 1% BSA, 5% normal horse serum (NHS; Cat#S-2000, Vector Laboratories), and 1.5% M.O.M in TBS for 1 hour at RT. Primary antibody was diluted in a solution of 1% BSA, 5% NHS, and 0.3% Triton X-100 in TBS to yield a final concentration of 10 μg/ml anti-BDNF. Sections were incubated for 2 nights at 4°C on a rotator. Then sections were incubated in biotinylated horse anti-mouse IgG secondary antibody (1:500; Cat#BA-2000, Vector Laboratories) in 1% BSA, and 0.3% Triton X-100 in TBS for 3 hours at RT, followed by Avidin-Biotin-Complex (ABC; ABC Elite kit; 1:1,000; Cat#PK-6100, Vector Laboratories) in 1% BSA in TBS for 1 hour at RT. Sections were washed in TBS and then Tris buffer (TB). Next, sections were reacted with 3, 3′-diaminobenzidine (DAB; Cat#DS905, Millipore-Sigma; 50 mg/100 ml in 0.1 M TB) in 40 μg/ml ammonium chloride (NH4Cl; Cat#A4514, Millipore-Sigma), 2 mg/ml D(+)–glucose (Cat#G5767, Millipore-Sigma), 10 mM nickel chloride (NiCl2; Cat#N6136, Millipore-Sigma), 3 μg/ml glucose oxidase (Cat#G2133-50KU, Millipore-Sigma), and then washed in TB. Labeled sections were mounted on gelatin-coated slides and dried overnight at RT. On the next day, sections were dehydrated with increasing concentrations of ethanol (70%, 2.5 minutes; 95%, 2.5 minutes; 100%, 5 minutes), cleared in Xylene (4 minutes; Cat#534056, Millipore-Sigma), and coverslipped with Permount (Cat#17986-01; Electron Microscopy Sciences).

Sections were examined using a microscope with brightfield and fluorescence detection (BX61, Olympus). Sections were photographed using a digital camera (Lumenera 3.0, Teledyne Instruments), images acquired using Infinity Analyze (Teledyne), and figures composed in Photoshop (v. 27; Adobe).

#### BDNF and GluR2/3 double staining

To distinguish BDNF protein expression in MCs, BDNF was double labeled with a marker of glutamatergic neurons, an antibody to the glutamate receptor 2/3 (GluR2/3, rabbit polyclonal, Cat#AB1506, Millipore-Sigma). The GluR2/3 antibody was raised against a synthetic peptide corresponding to the C-terminus of rat GluR2, which is almost identical to the C-terminus of rat GluR3. The antibody recognized a 110 kDa protein on Western blot corresponding to the molecular weight of GluR2 according to documentation of the manufacturer. In the past, GluR2/3 was shown to be an excellent marker of MCs, and the antibody has been well validated [76,77].

Free-floating sections were first washed in TB. Afterwards, sections were incubated in 0.25% Triton X-100 in TB (called TrisA below) and subsequently in 1% BSA and 0.25% Triton X-100 in TB (called TrisB below), for 10 minutes each. Then sections were blocked in 10% normal goat serum (NGS; Cat#S-1000, Vector Laboratories) and 3% DS, 1% BSA and 0.25% Triton X-100 in TB for 1 hour at RT. Sections were incubated overnight at 4°C on a rotator (Belly Dancer, Stovall) in primary antibodies (anti-BDNF, as described above for BDNF immunofluorescence, and 1:100 anti-GluR2/3) diluted in TrisB. On the following day, sections were rinsed in TrisA and subsequently in TrisB for 10 minutes each, then incubated in secondary antibodies, donkey anti–mouse Alexa Fluor 488 (described above) to allow BDNF-expressing neurons to fluorescence green and goat anti-rabbit Alexa Fluor 568 (1:500, Cat#811036, Invitrogen) to make GluR2/3-expressing neurons fluoresce red. Secondary antibodies were diluted in TrisB and incubated for 60 min at RT in the dark. Sections were washed in TB (3×5 min), mounted on gelatin-coated slides, and coverslipped with fluorescent mounting medium (Citifluor AF1;(Cat#17970-25, Electron Microscopy Sciences). Sections were examined using a laser scanning confocal microscope (LSM 880, Zeiss) and Zen 3.0 software (Zeiss). For z-stacks to discriminate co-localized BDNF and GluR2/3, images were acquired at 16-bit depth with a frame size of 2048 × 2048 pixels. Z-stack step size was 4 µm. The same parameters, including pixel time, gain and offset, were used for every image acquisition. Zen 3.2 Blue Edition software (Zeiss) was used offline to convert raw Zeiss image files (CZI format) into TIFF format for figures.

#### c-Fos and calretinin double staining

The procedures for double labeling with c-Fos and calretinin were the same as BDNF and GluR2/3 double staining but the primary antibodies were a rabbit monoclonal anti-c-Fos antibody (Cat#Ab190289, Abcam), and a mouse anti-calretinin antibody (Cat#6B3, Swant). The c-Fos antibody is raised against a recombinant full-length protein corresponding to human FOS and has been used extensively, with over 3000 peer-reviewed articles listed on the manufacturer’s website, and often in the mouse DG or cortex [78,79] and in our own work [80,81]. The calretinin antibody is raised against an epitope close to amino acids 125-178 which includes the linker region between EF-hand domains III and IV. It recognizes both calretinin-22k and full-length calretinin [82,83]. It also has been widely used and validated [84].

#### Estrogen receptors (ERs) and SATB1 double staining

The ERs expressed in MCs were studied by double immunofluorescence. MCs were identified by a mouse polyclonal antibody to a special AT-rich sequence-binding protein 1 (SATB1; 1:1000, #SC-376096, C6, Santa Cruz), a transcriptional factor that is expressed in many neuronal types [85] and was recently shown to be an excellent marker of MCs [86]. The blocking serum was 3% DS and 0.3% Triton X-100 in TBS and the secondary antibody was donkey anti-mouse Alexa Fluor 488.

For ERs, antibodies were used that detected ERα, ERβ, or GPER1. For ERα, a rabbit monoclonal anti-ERα antibody (1:2000, #04-820, clone 60C; Millipore-Sigma) was used that was raised to a KLH-conjugated synthetic peptide corresponding to amino acids 13-32 (C-ALLHQIQGNELEPLNRPQLK) of human ERα, which is the same in mouse. For ERβ, a rabbit polyclonal anti-ERβ (1:1000; Cat#PA1310B, Thermo Fisher) was used that has been commonly employed to study ERβ in the brain [87–89]. It was raised against a synthetic peptide corresponding to residues C(467) SSTEDSKNKESSQNLQSQ(485) of rat ERβ. The specificity of this antibody was demonstrated in western blot analysis of MCF7 and T47D cells, as well as LC-MS (https://www.thermofisher.com/antibody/product/Estrogen-Receptor-beta-Antibody-Polyclonal/PA1-310B), and specificity was also demonstrated by lack of detectable signal in a CRISPR-Cas9 knockout line [90]. For GPER1, a rabbit polyclonal anti-GPER1 antibody (1:1000, Cat#AB39742, Abcam) was used that has been previously validated by Western blot of brain tissues [91–95]. The immunogen used to generate the antibody could not be obtained because it is proprietary information. Staining was abolished upon omission of the primary antibody according to the manufacturer.

#### In situ hybridization for BDNF with immunofluorescence for MC markers

Sections were washed in 0.1M PB (3×5 minutes), mounted, and allowed to dry at RT overnight. The next day they were dehydrated (5 minutes, 50% EtOH, 10 minutes, 70% EtOH, 2×5 minutes, 100% EtOH) and then washed (5 minutes, ddH_2_O). A hydrophobic barrier was drawn around the sections with a pen provided by the manufacturer and allowed to dry for 5 minutes. Sections were then exposed to hydrogen peroxide solution provided by the manufacturer (RNAscope hydrogen peroxide solution; Biotechne) by placing drops on sections until the sections were covered and maintaining the slides in a horizontal position on a tray. Then they were washed in ddH_2_O in a 200 ml container and placed on a rotator to mix the solution (2×5 minutes). Excess solution was removed by tipping the slides. The next step was incubation in RNAscope target retrieval reagent for 15 minutes in a 200 ml container at 100°C. After washing (2×5 minutes, ddH_2_O, on a rotator), sections were washed for 5 minutes in RNAscope wash buffer. Next, slides were exposed to RNAscope manual pretreat reagent for 30 minutes in a 40°C water bath. Throughout this time sections were maintained in a horizontal position in the container described above to protect them from the water. Next, slides were washed 2×2 minutes with ddH_2_O, on a rotator, followed by 5 minutes in wash buffer. For the next step, drops of RNAscope Mm BDNF C1 probe were placed on sections and slides were incubated for 2 hours in the 40°C water bath. Unless specified otherwise, all washes below were 2×2 minutes in RNAscope wash buffer at RT. After washes, sections were exposed by drops to RNAscope hybridize AMP1 for 30 minutes in the 40°C water bath and washed again. Then RNAscope hybridize AMP2 was applied to sections and left for 30 minutes in the 40°C water bath followed by washes. Next, RNAscope hybridize AMP3 was used for 30 minutes in the 40°C water bath and then sections were washed. Afterwards, sections were immersed in RNAscope HRP C1 signal for 15 minutes in the 40°C water bath and washed. Sections were covered by drops of RNAscope Opal dye 520 (1:1000) in RNAscope TSA buffer in the 40°C water bath and washed. Then sections were incubated with drops of RNAscope HRP blocker in the 40°C water bath and washed. Next, immunostaining was initiated. For the following steps the slides were maintained horizontally in the container that had been used for the water bath before. Sections were exposed to drops of 3% NGS in TrisB for 1 hour at RT, then primary antibody (1:500 SATB1 or 1:100 Glur2/3, as described above) in 3% NGS in TrisB for 2 days at 4°C. They were washed (3×5 minutes in TB) and then incubated in secondary antibody (1:500, goat anti-mouse or goat anti-rabbit Alexa Fluor 568 as described above) diluted in TrisB for 1 hour at RT. After washes (3×5 minutes in TB) slides were placed in a refrigerator at 4°C overnight in a closed container with a damp cloth at the bottom. The next day they were left at RT for a few hours and then coverslipped in Citifluor AF1..

### V. Quantification

Quantification was blinded to sex and cycle stage. It was not possible to be blinded to the septotemporal level because the shapes of dorsal and ventral DG are very different. The hilus was defined as zone 4 of Amaral [96] and discriminated from CA3 and the GC layer by their distinct background staining.

#### Hilar cells with BDNF mRNA and the MC marker SATB1

Cells with BDNF mRNA (green) and/or SATB1+ (red) staining were counted manually using z-stacks from a dorsal and a ventral section selected at similar septotemporal levels (Dorsal: −1.9 mm from Bregma, 1.7 mm Interaural, www.mbl.org/atlas232; Ventral: −6.2 mm from Bregma, 3.8 mm Interaural; www.mbl.org/atlas170). Z-stacks were made with 4-5 focal planes through each section. Four-five planes were selected because this number sampled the top, center and bottom of most neurons. Step size was 4 µm. Cells were identified as double labeled if the cell had both BDNF mRNA (green) and SATB1 (red) expression in multiple focal planes of the cell. For cells that did not, they were defined as BDNF mRNA+ if there was BDNF mRNA labeling in the green channel and no detectable SATB1 staining in the red channel, and the opposite for SATB1+ cells. Labeling was defined as significantly greater fluorescence relative to background. To determine what was significantly greater fluorescence, z-stacks were converted to tiff files and imported into ImageJ where the intensity of fluorescence in and outside a cell could be measured. Three measurements were made of small regions of interest inside the cell, and three outside the cell. A significantly greater intensity was defined as a mean intensity that was >2 standard deviations higher than the background mean. Because of the time-consuming nature of this approach, it was done for a subset of cells and tissue sections.

For comparisons of intense BDNF mRNA, cells were counted if the mRNA fluorescent particles covered most of the SATB1+ MC soma rather than a small area of the cell. We acknowledge that this assessment was qualitative.

#### BDNF immunofluorescence in the IML

BDNF immunofluorescence in the IML was measured using a region of interest (ROI) surrounding the IML. For this purpose, the fluorescent image was imported into ImageJ. Then the IML that was fluorescent was outlined at 10x. The immunofluorescence (IF) of the outer molecular layer of the same section, which lacked detectable fluorescence, was subtracted from IML IF. For each mouse, 3 sections through the dorsal hippocampus (300 μm apart) were averaged to provide a value for dorsal hippocampus. Likewise, 3 sections (300 μm apart) were averaged for ventral hippocampus.

#### ERs in MCs using the MC marker SATB1

For double labeling of cells with the MC marker SATB1 and antibodies to ERs, cells were counted from analysis of red, green and merged images. Double labeled cells required red, green and yellow in the merged image. Pilot studies did not find red cells on top of green cells or vice versa, so z-stacks were not employed. As described above, 4 sections through the dorsal and ventral hippocampus (300 μm apart) were averaged.

#### C-Fos-labeled hilar cells with the MC marker calretinin

Counts were made manually from z-stacks, with red corresponding to the nucleus (c-Fos) and green for the cytoplasm (calretinin). Only one section was selected from each of the dorsal and ventral hippocampus.

### VI. Statistics

Results are presented as mean ± standard error of the mean (SEM). The p criterion was 0.05. Tests were conducted using Prism (v. 10.6.1, GraphPad). For comparison of two groups, unpaired t-tests were used if data were parametric, which was determined by the Shapiro-Wilk’s normality test and Bartlett’s test for homogeneity of variance. When there was a failure of normality, Mann-Whitney *U* tests were conducted; Bartlett’s tests shown no significant heteroscedasticity for any of the comparisons. For more than two groups, ANOVAs were used followed by Tukey-Kramer post-hoc tests for multiple comparisons. If data were nonparametric, a Kruskal-Wallis test was done. For all tests, the sample size was the number of mice, not the number of sections. Some comparisons have different sample sizes than others because tissues from some animals were damaged during processing.

## RESULTS

### I. BDNF expression in MCs

We first addressed whether BDNF protein could be detected in MC somata. Figure 1A shows a representative section from a male mouse stained with anti-BDNF and anti-GluR2/3 antibodies to identify BDNF (green) and GluR2/3 neurons (red). At low power the hilus is dominated by BDNF protein (Figure 1A) which at higher power appears to be in some cells that are double labeled, suggesting they are MCs (Figure 1A, arrows). BDNF protein was also expressed in the hilus around MCs, and in stratum lucidum of area CA3, reflecting the high BDNF protein in MFs shown previously [15–17]. Using a confocal microscope to examine the hilus, z-stacks confirmed that some hilar cells labeled with anti-BDNF antibodies also were stained with antibodies to GluR2/3 (Figure 1B-C).

**Figure 1.**
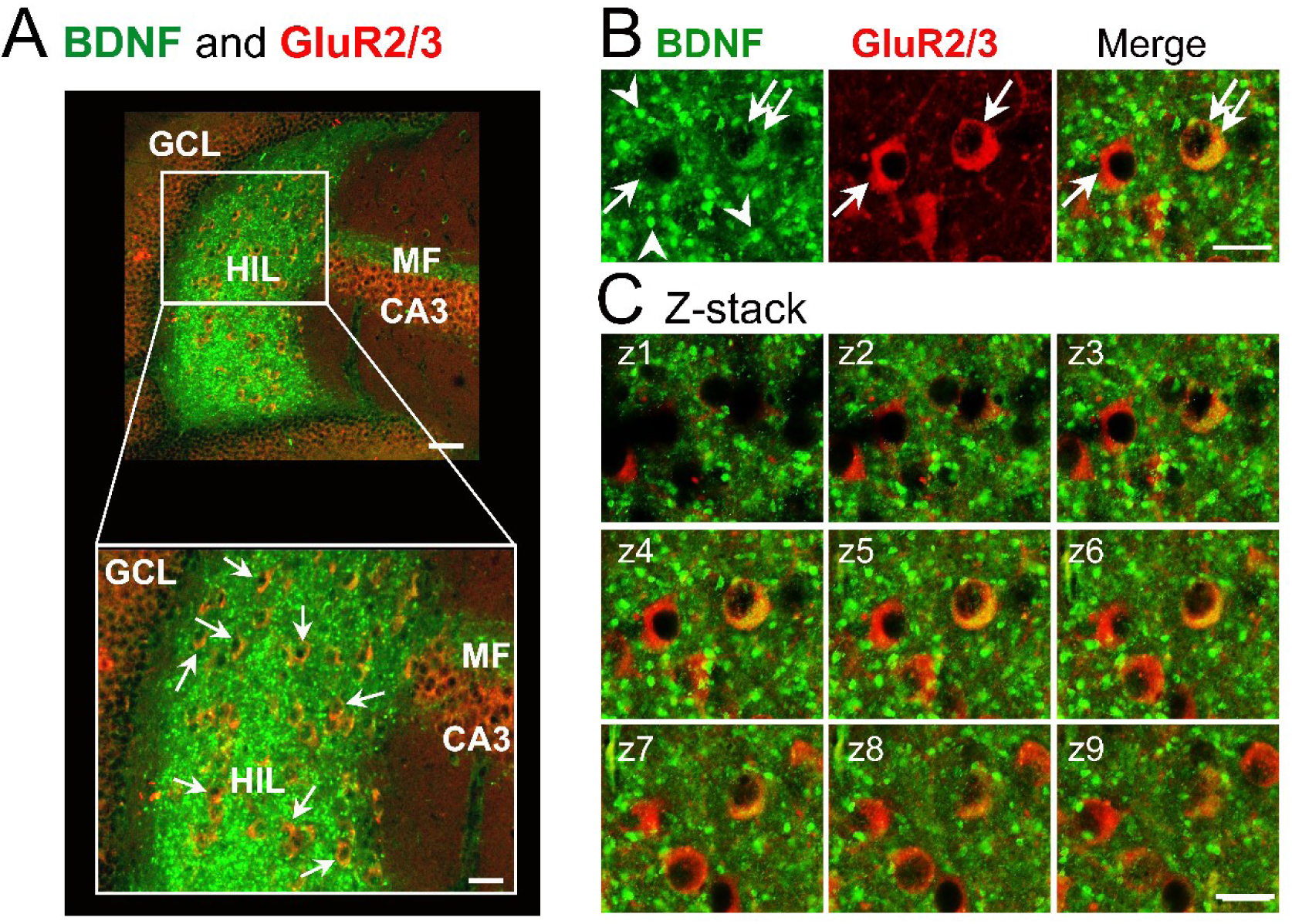
Robust BDNF protein expression in hilar mossy cell (MC) somata. **A.** Double labeling of BDNF protein (green) and GluR2/3 (red) in a horizontal section from an adult male mouse. Top: The granule cell layer (GCL) and CA3 pyramidal cell layer (CA3) express GluR2/3, whereas the mossy fiber (MF) axons of granule cells (GCs) express BDNF protein. In the hilus (HIL), there is both GluR2/3 and BDNF protein. Calibration, 25 µm. Bottom: The area surrounded by the box in the Top is expanded. White arrows point to double labeled hilar cells with both GluR2/3 and BDNF in the cytoplasm but not the nucleus, consistent with cytoplasmic expression of GluR2/3 and BDNF protein. Calibration, 15 µm. **B.** An area of the hilus in A is expanded to better illustrate double labeling. Left: The green channel shows a BDNF-labeled cell (double arrow) and a cell without BDNF protein (single arrow). There are numerous small profiles corresponding to axon terminals (arrowheads). Center: The red channel shows GluR2/3 labeled cells (arrows). Right: The merged image confirms one cell lacked BDNF protein (red; single arrow) whereas the other cell expressed BDNF protein and GluR2/3 (yellow/orange; double arrows). Calibration, 25 µm. **C.** A z stack shows sequential focal planes (z1-z9) of the double labeled cell in B. Both markers were present throughout the cell body. Calibration, 25 µm.

Given the intensity of MF BDNF in the hilus, it was difficult to quantify the number of MC somata that expressed BDNF protein. Therefore, we used in situ hybridization with probes for BDNF and double labeled the sections with the MC marker SATB1. We did not select GluR2/3 as a MC marker because pilot studies showed that it was incompatible with the protocol for in situ hybridization. Figure 2A-C shows the outcome, suggesting that BDNF mRNA was highly expressed in SATB1+ neurons.

**Figure 2.**
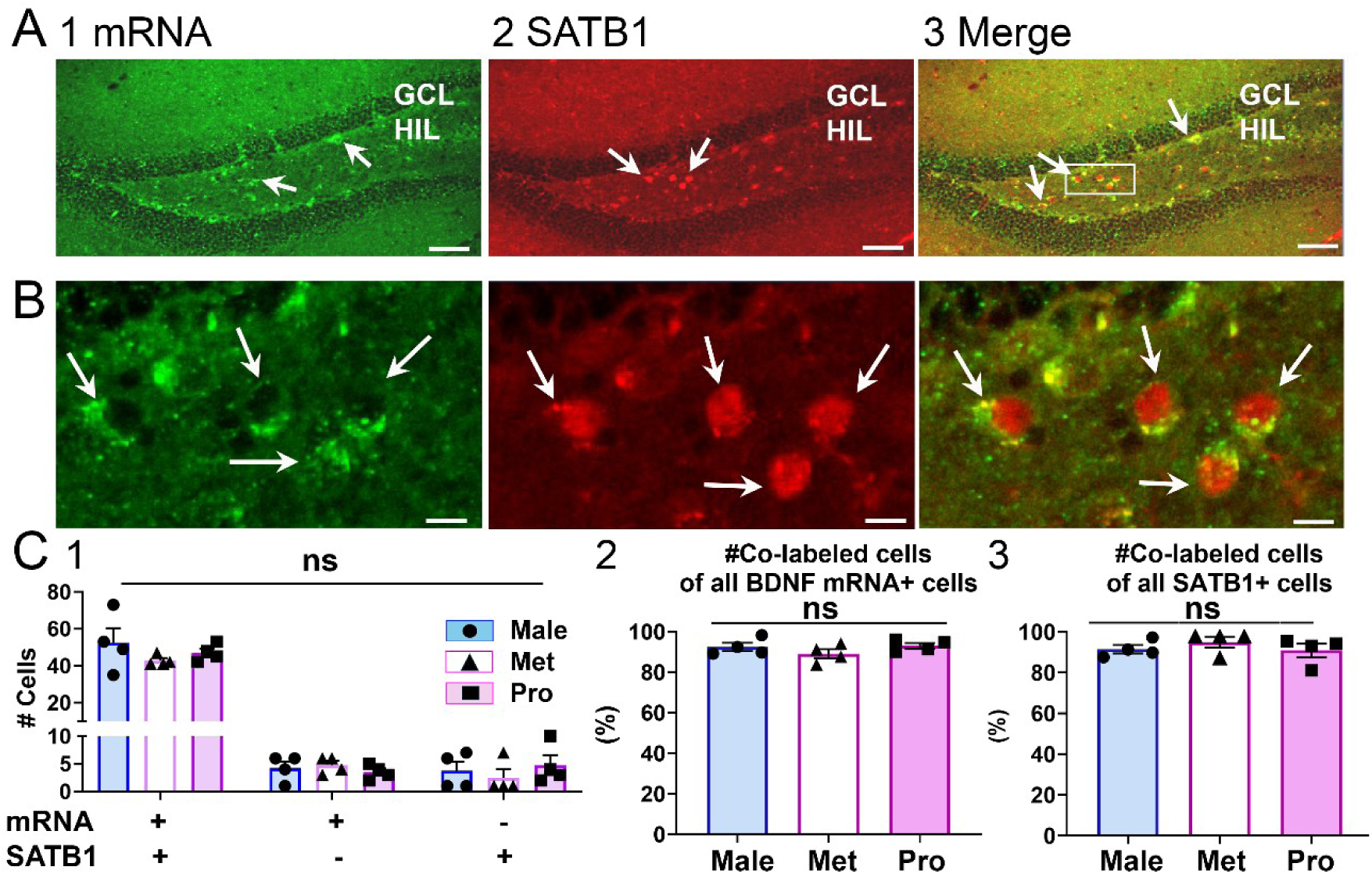
Robust BDNF mRNA in MCs. **A.** A section is shown from a male mouse that was stained for BDNF mRNA followed by immunofluorescence with an antibody to the MC marker SATB1. 1. Numerous cells in the hilus (HIL) have punctate staining in their cytoplasm, reflecting BDNF mRNA (green, arrows). GCL, granule cell layer. Calibration, 100 μm. 2. The red channel is shown to illustrate that numerous hilar somata express SATB1, indicating they are MCs (arrows). Calibration, 100 μm. 3. A merged image shows that numerous cells are double labeled (arrows), suggesting that many MCs have BDNF mRNA. Calibration, 100 μm. **B.** The area of A3 that is surrounded by a white box is expanded to better illustrate double labeled cells (arrows). Calibration, 20 μm. **C.** Quantification. 1. The number of hilar cells that expressed BDNF mRNA only, SATB1 only, or both BDNF mRNA and SATB1 were determined. A two-way ANOVA with type of labeling (e.g., BDNF mRNA only) and endocrine condition (e.g., male) as factors showed a significant effect of type of labeling (F(2,18)=216.50, p<0.0001) reflecting that most cells were double labeled, but no effect of endocrine condition (F(2,9)=1.02, p=0.400). 2. The number of double labeled cells is expressed as a percent of all hilar cells with BDNF mRNA. There were no significant differences (one-way ANOVA, F(2,9)=1.17, p=0.352, n=4/group). 3. The number of double labeled cells is expressed as a percent of all SATB1+ hilar cells. There were no significant differences (one-way ANOVA, F(2,9)=0.66, p=0.540, n=4/group).

After quantification (Figure 2C), there were few SATB1+ hilar neurons without BDNF mRNA (Figure 2C1). Instead, most SATB1+ hilar cells also expressed BDNF mRNA (Figure 2C1). Moreover, the percent of hilar cells expressing BDNF mRNA that also expressed SATB1 approached 100% (Figure 2C2), and the same was true for the percentage of SATB1+ hilar cells expressing BDNF mRNA (Figure 2C3). Results did not differ between males and females that were perfused on metestrous morning or proestrous morning (Figure 2C1-3). These data suggest that most MCs express BDNF mRNA and this occurs in both sexes. In addition, few hilar cells express BDNF mRNA besides MCs (Figure 2C1-3).

### II. BDNF protein in MC axons and its variation along the septotemporal axis

Having established BDNF mRNA was expressed in the majority of MC somata, we studied the MC axon plexus in the IML and found that BDNF-immunofluorescence was robust (Figure 3A1). Similar results were obtained for brightfield images using DAB (Figure 3A2). Although we do not have definitive proof that the IML BDNF protein is from MC axons, the only other neuronal types that innervate the IML do not make as dense an axon plexus in the IML as MCs. For example, the neurons of the lateral and medial supramammillary nucleus express BDNF protein but relatively little according to past studies [15,16]. The neurons of the medial septum and diagonal band project to the IML [97,98] but also showed few BDNF immunoreactive cells in past studies [15,16]. It is also notable that IML BDNF is likely to be due to anterograde transport, given the strong anterograde transport of BDNF by GCs [15,16].

**Figure 3.**
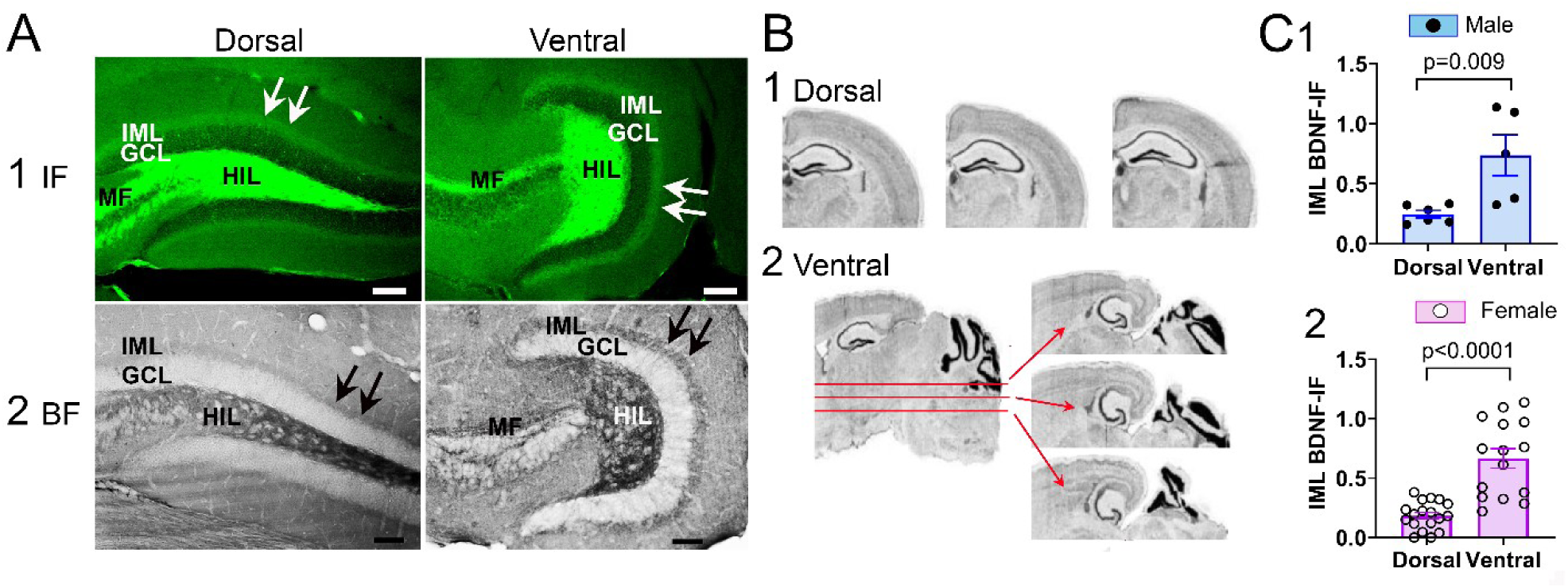
Robust BDNF protein expression in MC axons in the IML and its septotemporal variation. **A.** Examples of dorsal and ventral BDNF protein in the IML of male mice. 1. BDNF immunofluorescence (IF) in a male mouse is shown for a coronal section of dorsal DG (left) and horizontal section of ventral DG (right). Arrows point to the IML IF. GCL, granule cell layer; HIL, hilus; IML, inner molecular layer. Calibration, 100 µm. 2. BDNF protein expression in a different male mouse evaluated with brightfield (BF) microscopy also shows robust IML BDNF (arrows). Calibration, 100 µm. **B.** Methods for selecting dorsal and ventral sections. 1. Dorsal sections were sampled from one hemisphere cut in the coronal plane. Sections were selected approximately 300 μm apart as shown. 2. Ventral sections were sampled by cutting the second hemisphere horizontally. Sections were also approximately 300 μm apart. **C.** Greater BDNF protein expression in the ventral relative to the dorsal IML. 1. For male mice, the ventral IML had the greatest IF (Mann-Whitney *U* test, *U*=1.00, p=0.009, n=6 mice for dorsal data, n=5 for ventral). 2. Like males, female mice showed more ventral than dorsal IML BDNF-IF (unpaired t-test, t=5.04, df=20, p<0.0001). Metestrous females and proestrous females were pooled.

Next, we examined BDNF across the septotemporal axis. For this purpose, we sectioned one hemisphere in the coronal plane and selected dorsal sections and the other hemisphere was cut in the horizontal plane for ventral sections (see Methods; Figure 3B1-2). In 5 male mice we found that there was more BDNF protein in ventral than dorsal IML (Figure 3C1). These data suggest that dorsal MCs, which are primarily responsible for the ventral IML axons (Supplemental Figure 1), synthesize more BDNF than ventral MCs or transport it more to the ventral IML. Like males, females also showed more ventral IML BDNF immunofluorescence compared to the dorsal IML (Figure 3C2).

### III. BDNF protein in MC axons in the IML varies according to sex and estrous cycle stage

To evaluate males and females further, comparisons were made of males, females on metestrous morning, and females on proestrous morning. The results showed that metestrous females exhibited significantly less BDNF in the dorsal IML compared to males and proestrous females but males and proestrous females were not significantly different (Figure 4A-B, D). These data confirm in mice past findings in rats showing that MF BDNF is weak on metestrous morning compared to proestrous morning [42].

**Figure 4.**
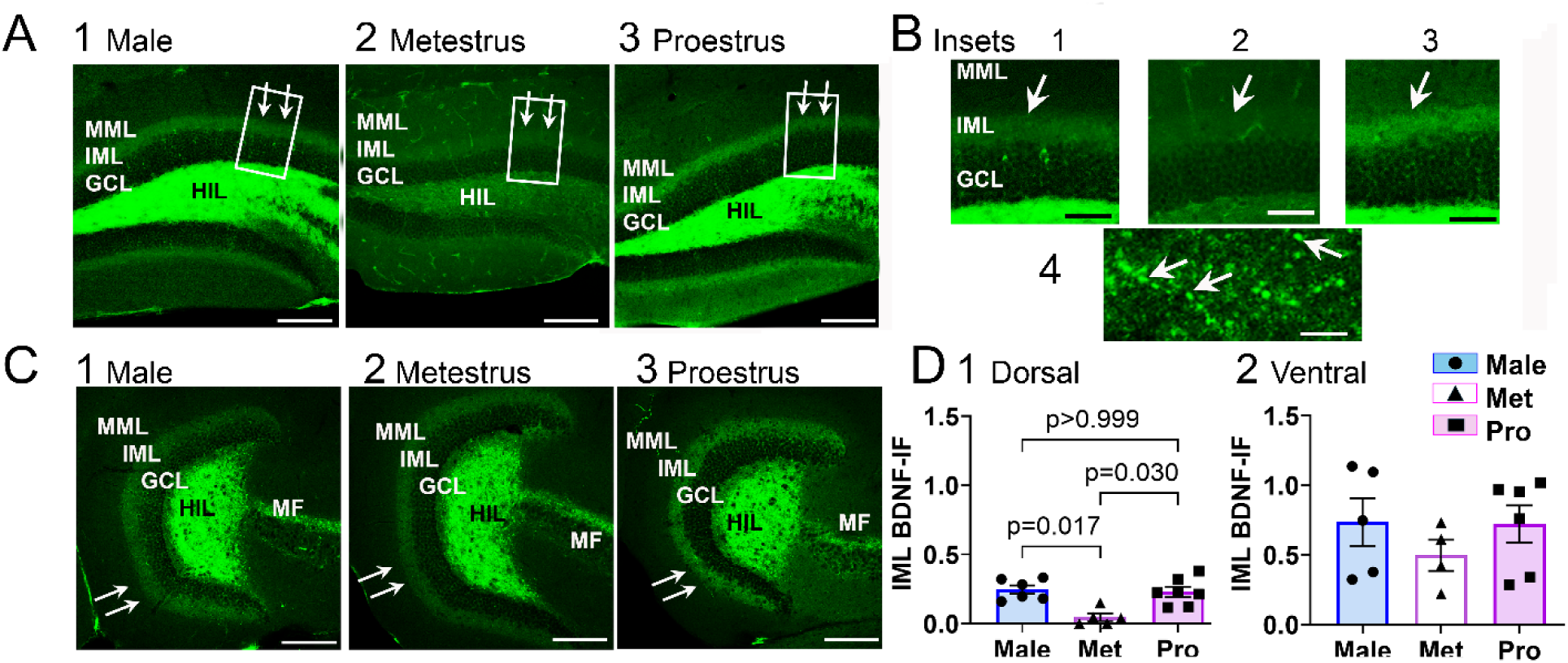
The influence of sex and the estrous cycle on IML BDNF-IF. **A.** Sex and the estrous cycle influence BDNF protein in the dorsal IML 1-3. Dorsal IML BDNF IF is compared for a male (1), metestrous female (2) and proestrous female (3). Sections were processed together and imaged with the same confocal settings. Arrows point to the IML. MML, middle molecular layer; IML, inner molecular layer; GCL, granule cell layer; HIL, hilus; MF, mossy fibers. Calibration, 200 μm. 1-3. The insets in A are expanded. Calibration, 50 μm. B. A high power image of the MC axon in the IML. Arrows point to axon terminals. Calibration, 15 μm. **C.** In contrast to the dorsal IML, sex and the estrous cycle do not influence BDNF IF in the ventral IML. 1-3. BDNF protein expression in the ventral IML (arrows) is compared for a male (1), metestrous female (2), and proestrous female (3). Calibration, 200 μm. **D.** Quantitation of IML BDNF IF. 1. Comparison of dorsal BDNF IF in 6 males, 5 metestrous females (Met), and 7 proestrous (Pro) females showed significant differences (Kruskal Wallis test: Kruskal Wallis statistic=9.141, p=0.005). Dunn’s post-hoc test showed significantly more BDNF IF in males than metestrous females (p=0.017) and more BDNF IF in proestrous females than metestrous females (p=0.030) but males and proestrous females were not different (p>0.999). Thus, males and proestrous females had higher IF than metestrous females. 2. Comparison of ventral IML BDNF IF in 5 males, 4 metestrous females, and 6 proestrous females showed no significant differences (one-way ANOVA, F(2,12)=0.736, p=0.500).

In contrast to the dorsal DG, the ventral IML showed no significant differences in BDNF immunofluorescence in males, metestrous females and proestrous females (Figure 4C-D). These data suggest that ventral MCs, which are primarily responsible for the dorsal IML axons (Supplemental Figure 1)[29,32], alter BDNF protein in response to sex and the estrous cycle. However, dorsal MCs, which project primarily to the ventral IML (Supplemental Figure 1)[29,32], vary less in response to sex and the estrous cycle.

### IV. BDNF mRNA shows septotemporal and endocrine influences complementary to those found for BDNF protein

To provide support for the explanation that dorsal IML BDNF was greater in males and proestrous females than metestrous females because ventral BDNF was higher in MC somata, we asked if ventral MC BDNF mRNA was relatively high in males and proestrous females relative to metestrous females. However, cells that expressed both BDNF mRNA and SATB1 did not show detectable variations among males, metestrous females and proestrous females for dorsal or ventral MCs (Figure 5A1-2). On the other hand, some ventral MCs had very intense BDNF mRNA labeling (Figure 5B-C). When these cells were quantified, they were significantly greater in males and proestrous females than metestrous females (Figure 5D). Therefore, the numbers of MCs with BDNF mRNA did not appear to differ but the amount of BDNF mRNA appeared to be greater in ventral MCs of males and proestrous females compared to metestrous females.

**Figure 5.**
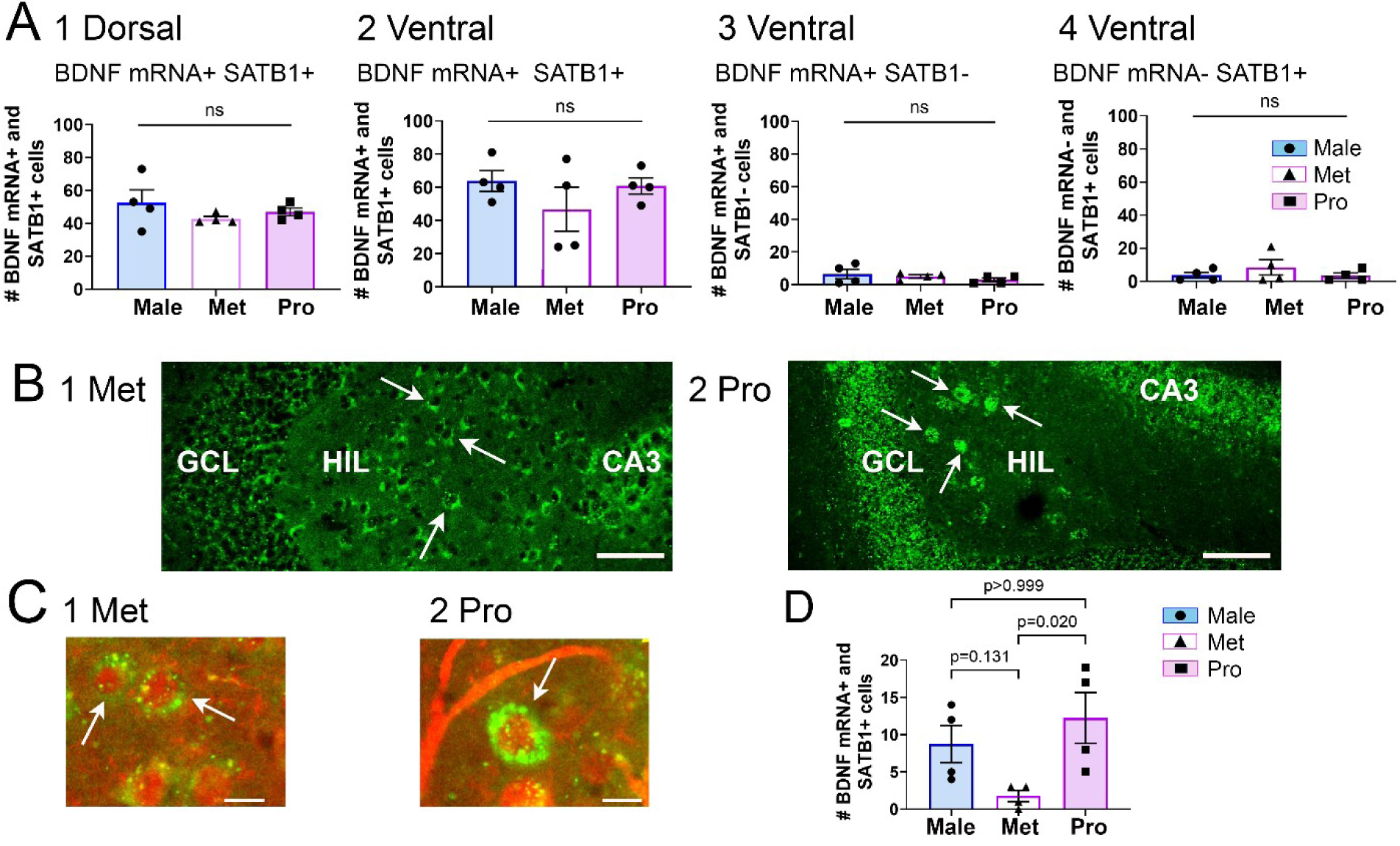
BDNF mRNA in ventral MCs shows intense labeling in males and proestrous females compared to metestrous females. **A.** Comparison of dorsal and ventral BDNF mRNA in hilar neurons. 1. Numerous neurons in the dorsal hilus (HIL) exhibited both BDNF mRNA and SATB1+ staining and there were no significant differences between males, metestrous females (Met), and proestrous females (Pro; one-way ANOVA, F(2,9)=1.04, p=0.393). The data in A are based on all BDNF mRNA+ cells, whereas only cells with intense labeling are examined in B-D. 2. Like dorsal hilus, there were no significant differences in double labeled cells in ventral hilus among males, metestrous females, and proestrous females (Kruskal Wallis test, Kruskal Wallis statistic 0.784, p=0.712). 3. In both dorsal and ventral hilus, few neurons expressed BDNF mRNA without SATB1 double labeling. The ventral data are shown (one-way ANOVA, F(2,9)=0.82, p=0.469). 4. In the dorsal and ventral hilus, few neurons expressed SATB1 protein without BDNF mRNA. The ventral data are shown (one-way ANOVA, F(2,9)=0.88, p=0.448). **B.** BDNF mRNA labeling is shown for the hilus of a female euthanized on metestrous morning (1) and proestrous morning (2). Arrows point to labeled neurons. Note intense labeling in some of the neurons from the proestrous female (arrows). GCL, granule cell layer, CA3, area CA3. Calibration, 100 µm. **C.** BDNF mRNA and SATB1 double labeled hilar cells (arrows) from a metestrous (1) and proestrous (2) female. Note intense BDNF mRNA labeling in the cell from the proestrous female. **D.** Quantification of double labeled cells that expressed strong BDNF mRNA labeling. There were significant differences (Kruskal Wallis test, Kruskal Wallis statistic, 7.911, p=0.005) with more cells with strong labeling on proestrous morning relative to metestrous morning (Dunn’s multiple comparisons post-hoc test, p=0.020), and no significant difference between males and proestrous females (p>0.999). There were no significant differences between males and metestrous females (p=0.131).

### V. Expression of ERs in MCs and the relevance to BDNF protein expression

Defining the types of ERs in MCs with MC markers has not been done. Therefore, we studied ERα, ERβ, and GPER1 in MCs. SATB1 was used as a marker of MCs because each of the antibodies to ERs were made in rabbit, and MC markers like GluR2/3 are also made in rabbit.

As shown in Figure 6A, ER expression was robust in SATB1-expressing hilar neurons. When quantified, a high percentage of SATB1+ neurons were also labeled with ER antibodies (Figure 6B). The percentages were robust in males, metestrous females and proestrous females (Figure 6B). When all 3 groups were pooled, there were more MCs labeled by GPER1- and ERβ compared to Erα (Figure 6C1). When the 3 groups were examined separately, GPER1-stained MCs outnumbered ERα-stained MCs on proestrus (Figure 6C2). These data indicate robust ER expression in MCs in both sexes and, when all groups were pooled, a higher percentage of MCs expressed GPER1 and ERβ relative to ERα. Moreover, GPER1-labeled MCs outnumbered ERα-expressing MCs on proestrous morning. The influence of sex and the estrous cycle on GPER1 expression was also shown in prior studies of the DG although MCs were not specifically examined [99].

**Figure 6.**
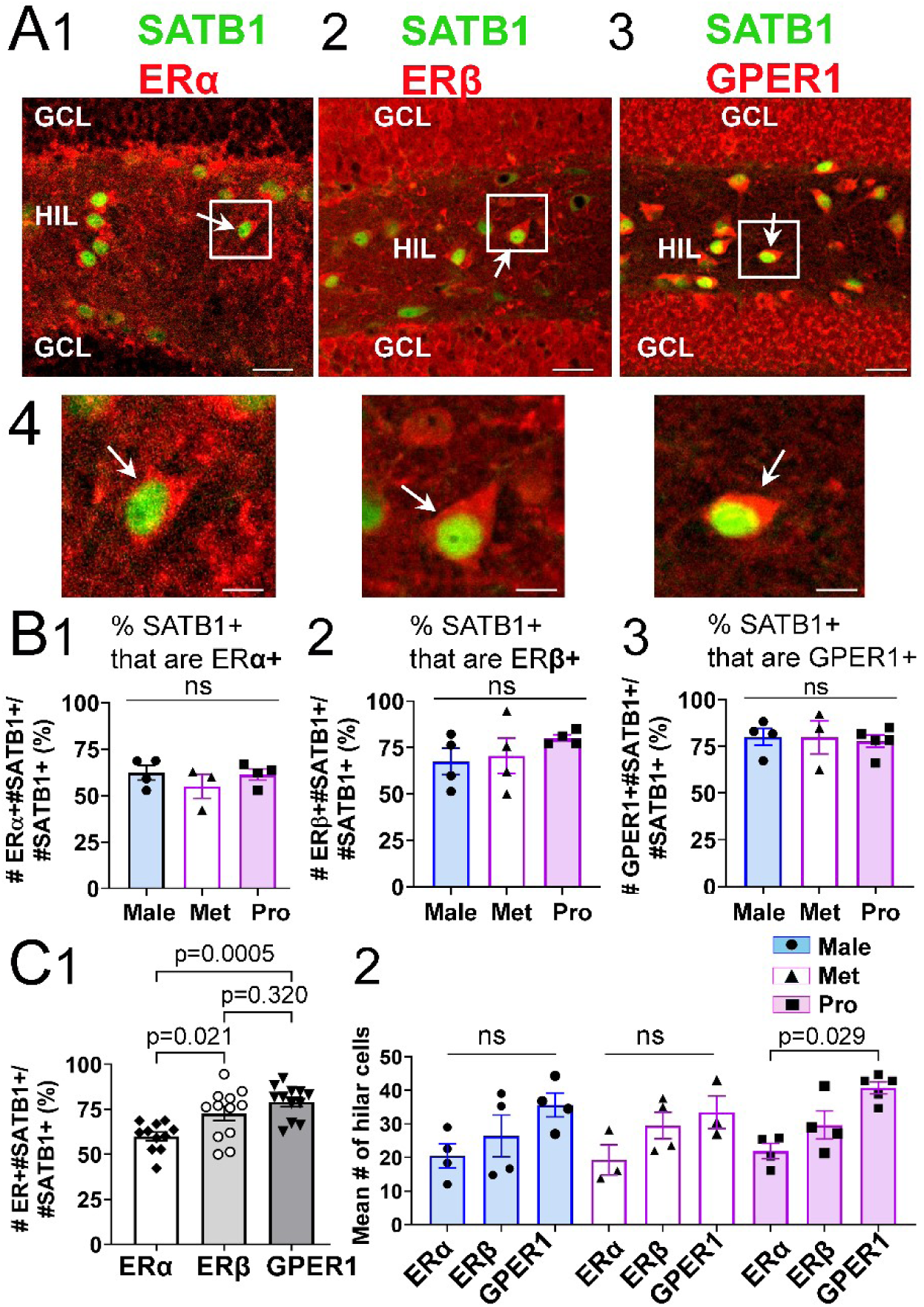
Robust expression of ERs in MCs. **A.** Sections were stained using an antibody to SATB1, a marker of MCs, and a second antibody to either ERα, ERβ, or GPER1. 1. Hilar cells expressing SATB1 (green) often co-expressed ERα (red; white arrow). This example is a dorsal section from a male. GCL, granule cell layer; HIL, hilus. Calibration, 50 μm. 2. Hilar cells expressing SATB1 (green) often co-expressed ERβ (red; arrow). This example is a dorsal section from a metestrous female. Calibration, 50 μm. 3. Hilar cells expressing SATB1 (green) often co-expressed GPER1 (red; arrow). The example is a dorsal section from a proestrous female. Calibration, 50 μm. 4. Left: The white box in A1 is expanded. Center: The white box in A2 is expanded. Right: The white box in A3 is expanded. Calibrations, 5 µm. **B.** Comparison of ERs in MCs for males, metestrous females (Met), and proestrous females (Pro). 1. A comparison of the mean # of double labeled hilar cells as a percent of all SATB1 cells shows the majority of SATB1 cells co-expressed ERα. There was no difference between males (n=4), females on metestrous morning (n=3), or females on proestrous morning (n=4; one-way ANOVA, F(2,8)=0.755, p=0.501). 2. A comparison of the mean # of double labeled hilar cells as a percent of all SATB1 cells shows the majority of SATB1 cells co-expressed ERβ but there was no influence of the endocrine condition (4 males, 4 metestrous females, 4 proestrous females; one-way ANOVA, F(2,9)=0.899, p=0.440). 3. A comparison of the mean # of double labeled hilar cells as a percent of all SATB1 cells shows that most SATB1 cells co-expressed GPER1 but there was no effect of the endocrine condition (4 males, 3 metestrous females, 5 proestrous females; one-way ANOVA, F(2,9)=0.064, p=0.938). **C.** Comparison of ERs in MCs for pooled data from males, metestrous females, and proestrous females. 1. The # of MCs that were ER+ (either ERα, ERβ, or GPER1) are expressed as a percent of all MCs, defined by SATB1. A one-way ANOVA showed a significant difference in ERs (one-way ANOVA, F(2,32)=9.37, p=0.0006) with more GPER1+ MCs than ERα+ MCs (p=0.0005) and more ERβ+ MCs than ERα+ MCs (p=0.021) but no difference between GPER1+ and ERβ+ MCs (p=0.320; Tukey-Kramer post-hoc tests). Same data as B. Pooled sample sizes: ERα, n=11; ERβ, n=12; GPER1, n=12. 2. Comparison of ERs in the 3 endocrine conditions (males, metestrous females, and proestrous females). A two-way ANOVA showed a significant effect of the type of ER (F(2,26)=11.93, p=0.0002) but no effect of endocrine condition (F(2,26)=0.737, p=0.488) with a significantly greater number of GPER1+ MCs than ERα+ MCs for proestrous females (Tukey-Kramer post-hoc test; p=0.029). Same data and sample sizes as B.

Previously, evidence was published that there is an εστρογεν ρεσπονσε-like element on the BDNF gene [45] and independently, 17β-estradiol can interact with the BDNF promoter to increase BDNF synthesis [46]. In light of these findings, and the high ER expression in MCs, it is possible that when 17β-estradiol rises in the serum during proestrous morning more BDNF is synthesized. Alternatively, ERs could use signaling pathways present in the MC cytoplasm to increase BDNF synthesis, such as the MAPK pathway [100–102].

### VI. Testing the activity-dependence of MC BDNF

Another potential explanation for the estrous cycle-dependence of BDNF protein in MCs is that rising levels of estradiol on proestrous morning increase activity of MCs, leading to activity-dependent BDNF synthesis [49,68]. To test this hypothesis, we examined MCs of females on proestrous and metestrous mornings using antibodies to a marker of recent neuronal activity, c-Fos, and the MC marker calretinin (Figure 7A-C). This approach was supported by prior studies showing that c-Fos protein is present in MCs that have increased excitability [81].

**Figure 7.**
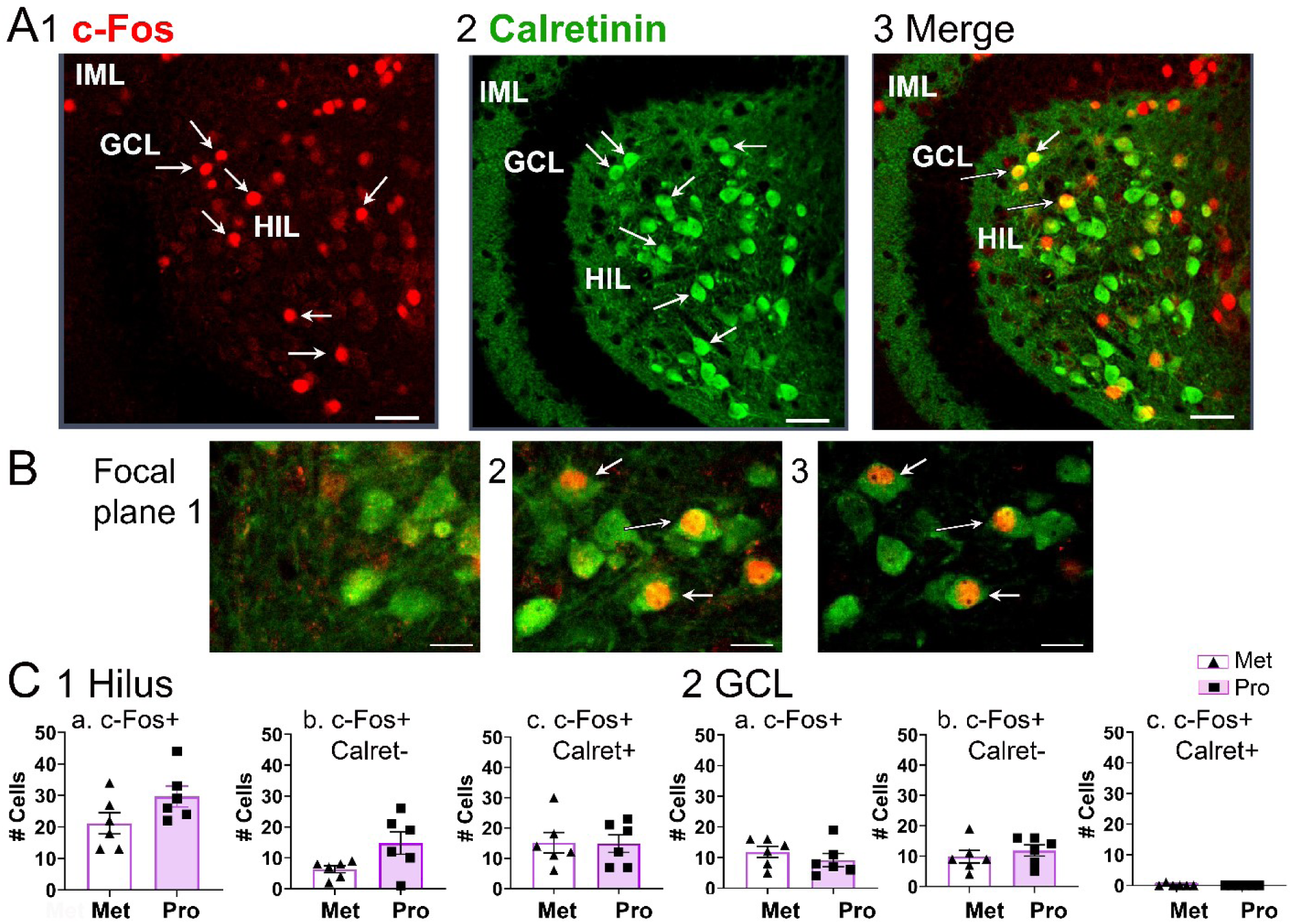
C-Fos protein in MCs on metestrous and proestrous mornings of the estrous cycle was not significantly different. **A.** A horizontal section from a proestrous female stained with antibodies to calretinin, a marker of MCs, and c-Fos, a marker of neuronal activity. Calibration, 50 μm. 1. c-Fos is expressed in numerous DG neurons, both in the granule cell layer (GCL) and hilus (HIL; arrows). Inner molecular layer, IML 2. Calretinin is expressed in many hilar neurons (arrows). 3. A merged image shows that some hilar neurons are double labeled (arrows). **A.** Sequential focal planes through the hilus of a section from a proestrous female. The arrows mark double labeled cells. 1. Double labeled cells have not yet come into focus. 2. Double labeling is coming into focus. 3. Double labeled cells are in focus. Calibration, 10 μm. 4. Comparison of hilar cells in metestrous females (Met) and proestrous females (Pro; n=6/group). For C1-2, there were no significant differences by unpaired t-tests as follows: a. C-Fos+ cells (t=1.81, df=10, p=0.101). b. C-Fos+/calretinin-cells (t=2.22, df=10, p=0.051). c. C-Fos+/calretinin+ cells (t=0.075, df=10, p=0.942). 5. GCL cells are compared for the same mice. a. C-Fos+ cells (t=0.936, df=10, p=0.372). b. C-Fos+/calretinin-cells (t=0.723, df=10, p=0.486). c. C-Fos+/calretinin+ cells ( t=1.00, df=10, p=0.341).

When metestrous females and proestrous females were compared, there was no significant difference in the numbers of MCs that expressed c-Fos (Figure 7C1). We also found that GC c-Fos was not significantly different between metestrous females and proestrous females (Figure 7C2). These data do not support the idea that BDNF in proestrous females is greater than metestrous females because MC activity is higher on proestrous morning.

In summary, ERs were robust in MCs, supporting the hypothesis that 17β-estradiol acting on MC ERs could increase BDNF synthesis. We did not find evidence that activity-dependent BDNF expression could explain why BDNF protein was greater in MC axons on proestrous relative to metestrous mornings.

## DISCUSSION

### I. Summary

The results demonstrate that MCs of the adult mouse show robust BDNF mRNA and protein in MC somata and there is strong BDNF protein expression in the IML. Furthermore, the majority of MCs express BDNF mRNA and protein. In contrast, other hilar neurons have much less BDNF mRNA. Interestingly, the ventral IML exhibited more BDNF protein than the dorsal IML. The results also show that BDNF protein in the dorsal IML varies depending on sex and estrous cycle stage, with more dorsal IML BDNF in males and females on proestrous morning compared to females on metestrous morning. In addition, ventral MC somata exhibited intense BDNF mRNA labeling on proestrous morning. Males also showed strong BDNF mRNA in ventral MCs but it was not significantly different from metestrous females. Since ventral MCs project primarily to the dorsal IML (Supplemental Figure 1)[29,32], the data suggest greater BDNF mRNA in ventral MC somata may explain greater dorsal IML BDNF.

One difference between proestrous and metestrous morning is that proestrous morning is the time during the estrous cycle when serum levels of 17β-estradiol peak [72,103]. Relatively low levels of serum 17β-estradiol are present on metestrous morning [72,103]. Therefore, higher serum levels of 17β-estradiol may stimulate MC BDNF synthesis on proestrous morning relative to metestrous morning. 17β-estradiol may stimulate MC BDNF synthesis by binding to MC ERs [45,46], so we determined if ERs were present in MCs. We found robust expression of all three ERs in the majority of MCs. We also studied c-Fos expression in MCs and found no significant difference between proestrous and metestrous mornings, suggesting activity-dependent BDNF synthesis [49,68] could not explain the relatively high levels of MC BDNF on proestrous mornings.

### II. BDNF expression in MCs

Prior studies have suggested that MCs express BDNF because the IML was shown to exhibit BDNF protein by immunohistochemistry [15] and might transported it anterogradely to the IML. Although anterograde transport of BDNF by MCs has not been shown, it is robust in GCs [15,16], so there is a precedent. Still, other BDNF expressing neurons project to the IML, such as neurons from the supramammillary nucleus [15]. Although BDNF expression in the supramammillary nucleus is s weak by immunohistochemistry [15], it is still a possible source of IML BDNF. Therefore, it was valuable to show that the majority of MC somata express BDNF using MC markers.Although this was difficult to show with immunohistochemistry because BDNF in GC axons overlaps MC somata, we could establish it using high magnification and BDNF mRNA. However, a limitation of this approach is that mRNA does not always translate into protein. For example, CA3 pyramidal cells have robust mRNA for BDNF but the GCs have greater BDNF protein than CA3 neurons [15].

### III. Septotemporal variation in MC BDNF

Males and females showed significantly greater BDNF protein expression in the IML of ventral DG relative to dorsal DG. A potential reason is based on the anatomical characteristics of the MC axon. Dorsal MCs project primarily to the ipsilateral ventral IML, rather than locally (Supplemental Figure 1)[29,32]. Therefore, the results presented here suggest that dorsal MCs, on average, synthesize more BDNF protein than ventral MCs. It is also possible that MCs of dorsal and ventral DG synthesize BDNF to the same extent, but dorsal MCs transport it more to their axons. The exception is the subset of MCs that show intense BDNF mRNA expression. They were located primarily in the ventral DG, giving them the potential to synthesize more BDNF. Since these intensely labeled MCs were primarily in proestrous females, they could contribute to the greater dorsal IML BDNF on proestrous morning.

The implications of the septotemporal variation in MC BDNF are interesting because they are relevant to the suggestion that dorsal DG is more involved in spatial memory and “cognitive” functions whereas ventral DG is more involved in contextual memory and “emotional” or “limbic” functions [37–41]. Consistent with this idea, overexpression of ventral BDNF in the hippocampus benefited mood and anxiety in a study of serotonin transporter knockout mice [104]. Another study suggested that BDNF contributes more to ventral DG functions than dorsal DG functions, but GC BDNF was the focus of that study, not MC BDNF [105]. It is a very relevant study, however, because it suggested that GC BDNF protein expression was greater in ventral DG than dorsal DG, reminiscent of the results we found for MCs. In addition, it was suggested that the role of ventral BDNF was mediated by immature adult-born neurons, which MCs innervate and nurture [106–108]. The influence of MCs on immature neurons may require BDNF since BDNF is critical to neuronal development [109–112].

It is notable that this is not the first time that differences in dorsal and ventral MCs have been noted. Prior studies show that gene expression is very different in dorsal and ventral MCs in rodents [36,113–115]. Dorsal and ventral MCs also differ in their electrophysiological properties [116], behavioral effects [117], the width of their projection to the IML [80,118], and their vulnerability in animal models of epilepsy [119]. Presumably the collective differences in dorsal and ventral MCs contribute to the dorsal and ventral differences in DG function.

### IV. Sex and estrous cycle variation in MC BDNF expression

Although there was increased expression of ventral BDNF in the IML compared to the dorsal DG, dorsal IML BDNF had more plasticity of expression in that it showed a sex and estrous cycle-dependent variation that the ventral IML did not. Thus, females that were examined on proestrous morning had significantly more dorsal IML BDNF than females on metestrous morning. Interestingly, there were no significant differences in dorsal IML BDNF between males and proestrous females although proestrous females had greater dorsal IML BDNF than metestrous females.

A caveat to these conclusions is that ventral IML BDNF showed a similar pattern as dorsal IML, with females on proestrous morning higher than metestrous morning, but differences were not significantly different.

### V. Estrogen and estrogen receptors in MCs

We questioned what might cause the sex and estrous cycle variation in dorsal IML BDNF and hypothesized that estrogen might increase BDNF since it has been suggested to increase BDNF synthesis. This hypothesis could be relevant to both males and females because hippocampal estrogen is present in both sexes [120–123]. However, serum levels of estrogen fluctuate in females during the estrous cycle, but it has not been shown that there are cyclic fluctuations in males. The mechanism by which estrogen induces BDNF synthesis may be that it binds to ERs that translocate to the nucleus and then act on an estrogen response-like element on the BDNF gene or interact with the BDNF promoter. Cytoplasmic ERs also can activate cytosolic signalling pathways that ultimately activate BDNF synthesis such as the mitogen activated protein kinase (MAPK) pathway [45,46]. An alternate hypothesis is that estrogen increases neuronal activity either by actions on GABAergic neurons [124], by influencing neuromodulators [125,126], or other mechanisms [127–129]. If estrogen increased MC neuronal activity, BDNF synthesis could increase because several BDNF transcripts are stimulated by activity [49,68].

To begin to address these hypotheses we first determined whether MCs have ERs. Although there is evidence that hilar neurons express ERs [50,52–55], results vary and past studies did not use MC markers so hilar GABAergic neurons were included. Our results showed that ERs were expressed in the majority of MCs, including ERα, ERβ and GPER1. ERβ and GPER1 labeling were high relative to ERα. When comparisons were made for each endocrine condition, there was significantly more GPER1 labeling of MCs on proestrous morning compared to ERα labeling at that cycle stage. Thus, there was a cycle and ER subtype-dependent variation in MC ERs.

These data suggest for the first time that most MCs express ERs and express all ERs. Furthermore, ERβ and GPER1 expression appeared to be particularly high in MCs. The results do not prove that estrogen action on MC ERs leads to increased BDNF synthesis but supports that potential mechanism. Regarding neuronal activity, we did not find evidence that MC activity is significantly different on proestrous morning compared to metestrous morning, suggesting that activity-dependent BDNF synthesis was not an explanation for the results. However, the studies of neuronal activity relied on c-Fos protein which is only one way to assess MC activity, and electrophysiological recordings of MCs might lead to a different result.

It is also important to note that some prior studies find few ERα- or ERβ-immunoreactive neurons in hippocampus, but most studies have focused on nuclear receptors. In our data, ERα and ERβ were typically cytoplasmic, suggesting they were membrane ERs.

### VI. BDNF in males

A prior study in rats showed that GC MFs exhibited lower BDNF protein in males than females on proestrous morning [42]. Therefore, it was surprising to find that MC BDNF was similar in male and proestrouss female mice.. One explanation is that there is a species difference. Another explanation is that regulation of GC MF BDNF differs from MC BDNF. It is also possible that there are differences between the prior study and the present study because of distinct antibodies to BDNF. In the past study in rats, an antibody was used that is no longer available. We selected a well-validated antibody for the present study that appears quite similar in its pattern of expression, but it still is a different antibody. Another point to note is that some of our data did suggest that male BDNF protein was weaker than females on proestrous morning. Thus, when comparing mean values, MCs on proestrous morning had the most MCs with intense BDNF mRNA. However, it was not significantly greater than males.

It is also important to consider whether 17β-estradiol levels in hippocampus are similar in males to those of proestrous females, so that there would be an equitable comparison of the effects of 17β-estradiol on BDNF levels. There are two main sources of 17β-estradiol: circulating levels of 17β-estradiol and 17β-estradiol synthesized in hippocampus [130]. However, results of past studies comparing hippocampal 17β-estradiol levels in males and female rodents vary [120–123], which could in part be due to different methods.

### VII. GC and MC BDNF

Our results suggest that MCs, like GCs, increase BDNF on proestrous morning relative to metestrous morning. Therefore, both the MC→GC synapse and the GC→PC synapse (the MF synapse) would have more BDNF on proestrous morning. In turn, synaptic plasticity may be enhanced in two pathways rather than just the MF pathway. Although little is known about the influence of BDNF on the MC→GC synapse, most hippocampal synapses are potentiated by BDNF, so it seems likely that BDNF release from MC terminals potentiates MC→GC synapses. If so, on proestrous morning there would be a potentially important increase in MC activation of GCs and subsequent GC activation of PCs. On the other hand, if BDNF release from MCs potentiates MC input to GABAergic interneurons, the excitation would be minimized. Future studies will be necessary to understand the role of MC BDNF on proestrous morning and whether it enhances GC activation of CA3.

### VIII. Limitations

There are several methodological limitations of this study. One is the ability of antibodies to accurately detect the protein of interest and RNAscope to accurately detect mRNA. However, the antibodies and RNAscope procedure have been used extensively and validated using diverse approaches [131]. One caveat is that the antibody to BDNF did not distinguish between proBDNF and BDNF.

Image acquisition can also have limitations. For example, during acquisition of images there needs to be consistent settings because otherwise one section can appear to be more labeled than the next. Therefore, our images were collected with the same settings (see Methods).

A potential limitation of defining cycle stage is the use of vaginal cytology. If only a few days are sampled, then one cannot be sure the mouse had regular estrous cycles, and one cannot predict cycle stage. For this reason, we sampled mice for at least 2 estrous cycles. We also have validated our methods in the past with hormone measurements and showed the predicted cycle stage based on vaginal cytology was the same as the stage demonstrated by hormone measurements [42].

Finally, it is important to bear in mind that the greater numbers of MCs with GPER1 and ERβ compared to ERα does not necessarily predict a greater effect of GPER1 and ERβ. One reason is that the ERs have different binding affinities for estradiol.

### IX. Conclusions

We conclude that MCs express BDNF in a robust manner in males and females, and as a result, could influence BDNF-dependent actions in the DG. Furthermore, during proestrous morning in females, greater ventral MC BDNF and greater dorsal IML BDNF could influence dorsal GCs more at that time. However, ventral GCs appear to have a greater and more stable input from BDNF-rich MC axons, potentially promoting ventral GC functions.

It has been suggested that dorsal GC functions are related to cognition, such as spatial memory. In light of this view, the data suggest that MC BDNF acting on dorsal GCs might allow more cognitive functions of GCs on proestrous morning. That idea would be consistent with the need of proestrous females to find a mate on proestrus, since rodents typically mate during the evening of proestrus [5,103]. The ability to find a mate may require greater spatial cognitive functions, which could be enhanced by greater MC BDNF input to dorsal GCs. In contrast, ventral GCs are considered to serve functions related to mood, anxiety, stress and contextual memory. These functions might benefit from robust MC BDNF at all times, not only on proestrous mornings.

Another conclusion is that MCs express all types of ERs in a robust manner and do so in both sexes and across both stages of the estrous cycle that were examined. Robust MC ERs place MCs in a strong position to mediate diverse effects of estrogen. Some of these effects overlap with those for BDNF, such as promoting plasticity of synaptic transmission, dendritic spines, and promoting adult neurogenesis [132]. Regarding spines, it is notable that spines are abundant on MCs and estrogen could be important to maintain these spines [129,133,134]. Other effects of estrogen could be to protect MCs in disease, since MCs are highly sensitive to insults and injury, and estrogen protects hilar neurons [135,136].

## Supplemental Figures

**Supplemental Figure 1.**
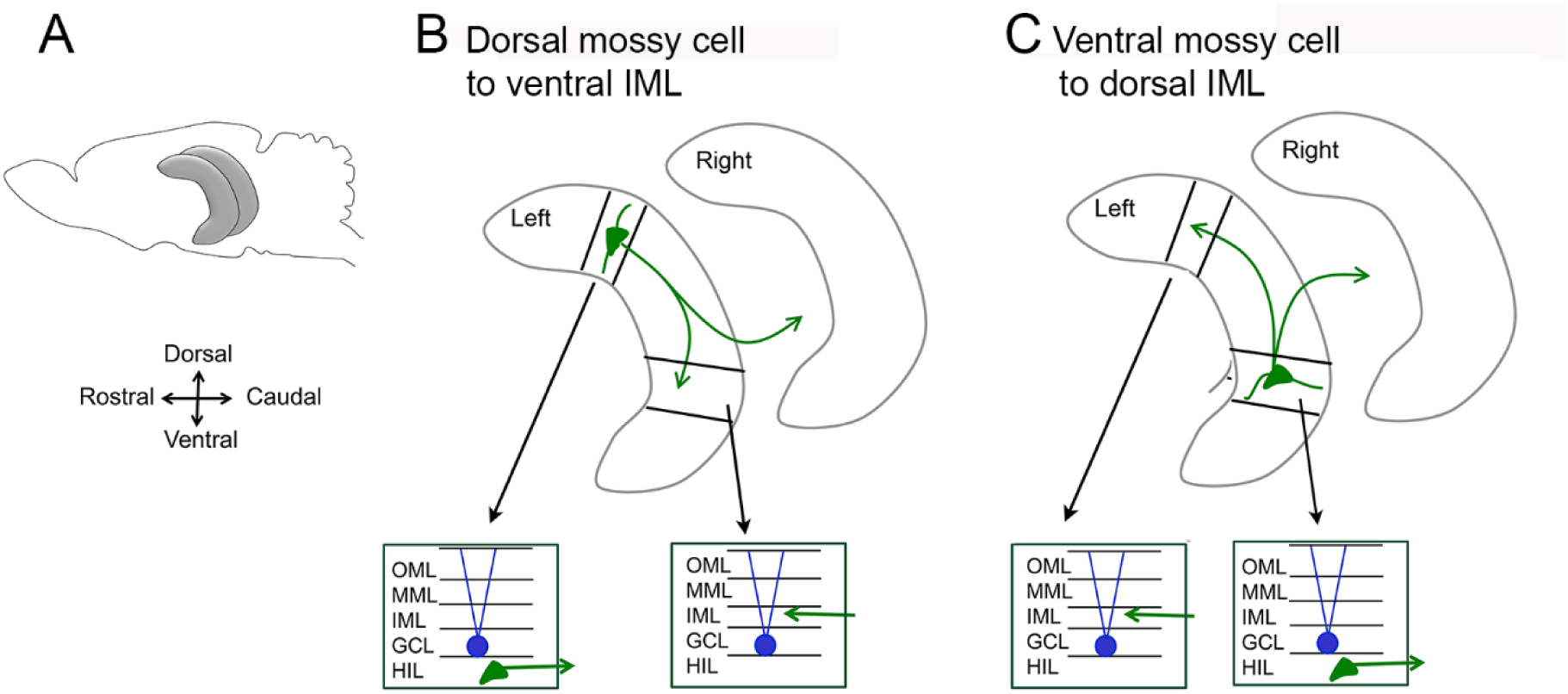
Dorsal and ventral MCs project mainly to distal locations. **A.** A sagittal view of the mouse brain shows the location of the two hippocampi (gray). Dorsal is up and rostral is to the left. **B.** Top: The left and right hippocampi are shown with the same orientation as in A. Within the left hippocampus is a dorsal MC soma (green shape) with lines as its dendrites and its axon indicated by green arrows. The main projection is distant to the MC soma in the molecular layer of the ventral DG. It also projects contralaterally to the molecular layer. Note that the MC axon also has collaterals near the cell body (not shown) that are in the hilus and molecular layer [28,31]. Bottom: The circuitry in the DG of the lamella near the dorsal MC cell body is shown at left and the circuitry of a ventral lamella is on the right. The lamella with the dorsal MC cell body shows the cell body is in the hilus and the main axon projection as a green arrow exiting the lamella. In the ventral lamella the termination of the distal axon projection is shown as a green arrow entering the IML. **C.** Diagrams analogous to those in B illustrate the cell body location and main axon projection of ventral MCs. In this case the cell body in the hilus is in the ventral lamella and in the dorsal lamella are the terminations of the ventral MC axons in the dorsal IML.

## Data Availability Statement

All data will be freely available at Open Science Framework (www.osf.io)

## Acknowledgments

We thank Meghan Kennedy for assistance and Drs. Neil MacLusky and Solomon Haizel for comments on the manuscript.

## Author Contributions

CC: Conceptualization, experimental design, data acquisition, analysis, figure preparation, writing, editing; MR: Data acquisition; FJ: Data acquisition; HS:

Conceptualization, experimental design, data analysis, figure preparation, writing, editing.

## REFERENCES

1. Chao MV and Hempstead BL. P75 and Trk: A two-receptor system. Trends Neurosci. 1995; 18: 321–6.

2. Barde YA. The physiopathology of brain-derived neurotrophic factor. Physiol Rev. 2025; 105: 2073–2140.

3. Danzer SC, Kotloski RJ, Walter C, Hughes M, and McNamara JO. Altered morphology of hippocampal dentate granule cell presynaptic and postsynaptic terminals following conditional deletion of TrkB. Hippocampus. 2008; 18: 668–78.

4. von Bohlen und Halbach O, Krause S, Medina D, Sciarretta C, Minichiello L, and Unsicker K. Regional- and age-dependent reduction in TrkB receptor expression in the hippocampus is associated with altered spine morphologies. Biol Psychiatry. 2006; 59: 793–800.

5. Adachi M, Barrot M, Autry AE, Theobald D, and Monteggia LM. Selective loss of brain-derived neurotrophic factor in the dentate gyrus attenuates antidepressant efficacy. Biol Psychiatry. 2008; 63: 642–9.

6. Li Y, Luikart BW, Birnbaum S, Chen J, Kwon CH, Kernie SG, Bassel-Duby R, and Parada LF. TrkB regulates hippocampal neurogenesis and governs sensitivity to antidepressive treatment. Neuron. 2008; 59: 399–412.

7. Ma DK, Jang MH, Guo JU, Kitabatake Y, Chang ML, Pow-Anpongkul N, Flavell RA, Lu B, Ming GL, and Song H. Neuronal activity-induced gadd45b promotes epigenetic DNA demethylation and adult neurogenesis. Science. 2009; 323: 1074–7.

8. Bergami M, Rimondini R, Santi S, Blum R, Gotz M, and Canossa M. Deletion of TrkB in adult progenitors alters newborn neuron integration into hippocampal circuits and increases anxiety-like behavior. Proc Natl Acad Sci U S A. 2008; 105: 15570–5.

9. Wang YJ, Wang YX, Zou CZ, Zhang WJ, Pan W, Wang JQ, Wang H, Qian X, Han GJ, Liu FG, and Wang J. PGC-1α promotes the survival of newborn neurons within AD hippocampus through activation of the FNDC5/BDNF/TrkB signaling pathway. Front Mol Neurosci. 2025; 18: 1688694.

10. Isono J, Nakagawasai O, Takahashi K, Nemoto W, and Tan-No K. Microinjection of brain-derived neurotrophic factor into the dorsal hippocampus alleviates cognitive dysfunction and increases hippocampal cell proliferation in olfactory bulbectomized mice. Neurosci Lett. 2025; 859-861: 138257.

11. Lee J, Duan W, and Mattson MP. Evidence that brain-derived neurotrophic factor is required for basal neurogenesis and mediates, in part, the enhancement of neurogenesis by dietary restriction in the hippocampus of adult mice. J Neurochem. 2002; 82: 1367–75.

12. Schabitz WR, Steigleder T, Cooper-Kuhn CM, Schwab S, Sommer C, Schneider A, and Kuhn HG. Intravenous brain-derived neurotrophic factor enhances poststroke sensorimotor recovery and stimulates neurogenesis. Stroke. 2007; 38: 2165–72.

13. Chan JP, Cordeira J, Calderon GA, Iyer LK, and Rios M. Depletion of central BDNF in mice impedes terminal differentiation of new granule neurons in the adult hippocampus. Mol Cell Neurosci. 2008; 39: 372–83.

14. Pinnock SB and Herbert J. Brain-derived neurotropic factor and neurogenesis in the adult rat dentate gyrus: Interactions with corticosterone. Eur J Neurosci. 2008; 27: 2493–500.

15. Conner JM, Lauterborn JC, Yan Q, Gall CM, and Varon S. Distribution of brain-derived neurotrophic factor (BDNF) protein and mRNA in the normal adult rat cns: Evidence for anterograde axonal transport. J Neurosci. 1997; 17: 2295–313.

16. Yan Q, Rosenfeld RD, Matheson CR, Hawkins N, Lopez OT, Bennett L, and Welcher AA. Expression of brain-derived neurotrophic factor protein in the adult rat central nervous system. Neuroscience. 1997; 78: 431–48.

17. Dieni S, Matsumoto T, Dekkers M, Rauskolb S, Ionescu MS, Deogracias R, Gundelfinger ED, Kojima M, Nestel S, Frotscher M, and Barde YA. BDNF and its pro-peptide are stored in presynaptic dense core vesicles in brain neurons. J Cell Biol. 2012; 196: 775–88.

18. Brigadski T and Lessmann V. The physiology of regulated BDNF release. Cell Tissue Res. 2020; 382: 15–45.

19. Schildt S, Endres T, Lessmann V, and Edelmann E. Acute and chronic interference with BDNF/TrkB-signaling impair LTP selectively at mossy fiber synapses in the CA3 region of mouse hippocampus. Neuropharmacology. 2013; 71: 247–54.

20. Schjetnan AG and Escobar ML. In vivo BDNF modulation of hippocampal mossy fiber plasticity induced by high frequency stimulation. Hippocampus. 2012; 22: 1–8.

21. Martinez-Moreno A, Rivera-Olvera A, and Escobar ML. BDNF induces in vivo long-lasting enhancement of synaptic transmission and structural reorganization at the hippocampal mossy fibers in a transcription and translation-independent manner. Neurobiol Learn Mem. 2020; 167: 107125.

22. Scharfman HE. Hyperexcitability in combined entorhinal/hippocampal slices of adult rat after exposure to brain-derived neurotrophic factor. J Neurophysiol. 1997; 78: 1082–95.

23. Myers CE and Scharfman HE. Pattern separation in the dentate gyrus: A role for the CA3 backprojection. Hippocampus. 2011; 21: 1190–215.

24. Rolls ET. The mechanisms for pattern completion and pattern separation in the hippocampus. Front Syst Neurosci. 2013; 7: 74.

25. Garad M, Edelmann E, and Lessmann V. Long-term depression at hippocampal mossy fiber-CA3 synapses involves BDNF but is not mediated by p75NTR signaling. Sci Rep. 2021; 11: 8535.

26. Messaoudi E, Ying SW, Kanhema T, Croll SD, and Bramham CR. Brain-derived neurotrophic factor triggers transcription-dependent, late phase long-term potentiation in vivo. J Neurosci. 2002; 22: 7453–61.

27. Olofsdotter K, Lindvall O, and Asztely F. Increased synaptic inhibition in dentate gyrus of mice with reduced levels of endogenous brain-derived neurotrophic factor. Neuroscience. 2000; 101: 531–9.

28. Chang H, Leem YH, Park J, Park JE, Yang SJ, and Kim HS. High-intensity interval training improves memory deficits in obese mice by enhancing osteocalcin-driven astrocytic BDNF expression and stimulating hippocampal neurogenesis. Neurochem Res. 2025; 50: 248.

29. Scharfman HE. The enigmatic mossy cell of the dentate gyrus. Nat Rev Neurosci. 2016; 17: 562–75.

30. Wenzel HJ, Buckmaster PS, Anderson NL, Wenzel ME, and Schwartzkroin PA. Ultrastructural localization of neurotransmitter immunoreactivity in mossy cell axons and their synaptic targets in the rat dentate gyrus. Hippocampus. 1997; 7: 559–70.

31. Buckmaster PS, Wenzel HJ, Kunkel DD, and Schwartzkroin PA. Axon arbors and synaptic connections of hippocampal mossy cells in the rat in vivo. J Comp Neurol. 1996; 366: 271–92.

32. Scharfman HE and Myers CE. Hilar mossy cells of the dentate gyrus: A historical perspective. Front Neural Circuits. 2012; 6: 106.

33. Nasrallah K, Frechou MA, Yoon YJ, Persaud S, Goncalves JT, and Castillo PE. Seizure-induced strengthening of a recurrent excitatory circuit in the dentate gyrus is proconvulsant. Proc Natl Acad Sci U S A. 2022; 119: e2201151119.

34. Nasrallah K, Berthoux C, Hashimotodani Y, Chavez AE, Gulfo MC, Lujan R, and Castillo PE. Retrograde adenosine a2a receptor signaling facilitates excitatory synaptic transmission and seizures. Cell Rep. 2024; 43: 114382.

35. Hashimotodani Y, Nasrallah K, Jensen KR, Chavez AE, Carrera D, and Castillo PE. LTP at hilar mossy cell-dentate granule cell synapses modulates dentate gyrus output by increasing excitation/inhibition balance. Neuron. 2017; 95: 928–943 e3.

36. Cembrowski MS, Wang L, Sugino K, Shields BC, and Spruston N. Hipposeq: A comprehensive RNA-seq database of gene expression in hippocampal principal neurons. Elife. 2016; 5: e14997.

37. Moser MB and Moser EI. Functional differentiation in the hippocampus. Hippocampus. 1998; 8: 608–19.

38. Bannerman DM, Sprengel R, Sanderson DJ, McHugh SB, Rawlins JN, Monyer H, and Seeburg PH. Hippocampal synaptic plasticity, spatial memory and anxiety. Nat Rev Neurosci. 2014; 15: 181–92.

39. Lee AR, Kim JH, Cho E, Kim M, and Park M. Dorsal and ventral hippocampus differentiate in functional pathways and differentially associate with neurological disease-related genes during postnatal development. Front Mol Neurosci. 2017; 10: 331.

40. Kheirbek MA, Drew LJ, Burghardt NS, Costantini DO, Tannenholz L, Ahmari SE, Zeng H, Fenton AA, and Hen R. Differential control of learning and anxiety along the dorsoventral axis of the dentate gyrus. Neuron. 2013; 77: 955–68.

41. Kheirbek MA and Hen R. Dorsal vs ventral hippocampal neurogenesis: Implications for cognition and mood. Neuropsychopharmacology. 2011; 36: 373–4.

42. Scharfman HE, Mercurio TC, Goodman JH, Wilson MA, and MacLusky NJ. Hippocampal excitability increases during the estrous cycle in the rat: A potential role for brain-derived neurotrophic factor. J Neurosci. 2003; 23: 11641–52.

43. Harte-Hargrove LC, Maclusky NJ, and Scharfman HE. Brain-derived neurotrophic factor-estrogen interactions in the hippocampal mossy fiber pathway: Implications for normal brain function and disease. Neuroscience. 2013; 239: 46–66.

44. Scharfman HE and MacLusky NJ. Estrogen and brain-derived neurotrophic factor (BDNF) in hippocampus: Complexity of steroid hormone-growth factor interactions in the adult cns. Front Neuroendocrinol. 2006; 27: 415–35.

45. Sohrabji F, Miranda RC, and Toran-Allerand CD. Identification of a putative estrogen response element in the gene encoding brain-derived neurotrophic factor. Proc Natl Acad Sci U S A. 1995; 92: 11110–4.

46. Deb P, Chini A, Guha P, Rishi A, Bhan A, Brady B, Perrotti LI, and Mandal SS. Dynamic regulation of BDNF gene expression by estradiol and lncRNA HOTAIR. Gene. 2024; 897: 148055.

47. Blurton-Jones M and Tuszynski MH. Estradiol-induced modulation of estrogen receptor-β and GABA within the adult neocortex: A potential transsynaptic mechanism for estrogen modulation of BDNF. J Comp Neurol. 2006; 499: 603–12.

48. Blurton-Jones M, Kuan PN, and Tuszynski MH. Anatomical evidence for transsynaptic influences of estrogen on brain-derived neurotrophic factor expression. J Comp Neurol. 2004; 468: 347–60.

49. Pattabiraman PP, Tropea D, Chiaruttini C, Tongiorgi E, Cattaneo A, and Domenici L. Neuronal activity regulates the developmental expression and subcellular localization of cortical BDNF mRNA isoforms in vivo. Mol Cell Neurosci. 2005; 28: 556–70.

50. Orikasa C, McEwen BS, Hayashi H, Sakuma Y, and Hayashi S. Estrogen receptor α, but not β, is expressed in the interneurons of the hippocampus in prepubertal rats: An in situ hybridization study. Dev Brain Res. 2000; 120: 245–54.

51. Hajszan T, Milner TA, and Leranth C. Sex steroids and the dentate gyrus. Prog Brain Res. 2007; 163: 399–415.

52. Milner TA, McEwen BS, Hayashi S, Li CJ, Reagan LP, and Alves SE. Ultrastructural evidence that hippocampal α estrogen receptors are located at extranuclear sites. J Comp Neurol. 2001; 429: 355–71.

53. Milner TA, Ayoola K, Drake CT, Herrick SP, Tabori NE, McEwen BS, Warrier S, and Alves SE. Ultrastructural localization of estrogen receptor β immunoreactivity in the rat hippocampal formation. J Comp Neurol. 2005; 491: 81–95.

54. Milner TA, Thompson LI, Wang G, Kievits JA, Martin E, Zhou P, McEwen BS, Pfaff DW, and Waters EM. Distribution of estrogen receptor β containing cells in the brains of bacterial artificial chromosome transgenic mice. Brain Res. 2010; 1351: 74–96.

55. Waters EM, Thompson LI, Patel P, Gonzales AD, Ye HZ, Filardo EJ, Clegg DJ, Gorecka J, Akama KT, McEwen BS, and Milner TA. G-protein-coupled estrogen receptor 1 is anatomically positioned to modulate synaptic plasticity in the mouse hippocampus. J Neurosci. 2015; 35: 2384–97.

56. Stone DJ, Rozovsky I, Morgan TE, Anderson CP, and Finch CE. Increased synaptic sprouting in response to estrogen via an apolipoprotein E-dependent mechanism: Implications for Alzheimer’s disease. J Neurosci. 1998; 18: 3180–5.

57. Scharfman HE. The role of nonprincipal cells in dentate gyrus excitability and its relevance to animal models of epilepsy and temporal lobe epilepsy. Adv Neurol. 1999; 79: 805–20.

58. Picazo O, Azcoitia I, and Garcia-Segura LM. Neuroprotective and neurotoxic effects of estrogens. Brain Res. 2003; 990: 20–7.

59. Veiga S, Garcia-Segura LM, and Azcoitia I. Neuroprotection by the steroids pregnenolone and dehydroepiandrosterone is mediated by the enzyme aromatase. J Neurobiol. 2003; 56: 398–406.

60. Azcoitia I, Sierra A, and Garcia-Segura LM. Estradiol prevents kainic acid-induced neuronal loss in the rat dentate gyrus. Neuroreport. 1998; 9: 3075–9.

61. Ervin KS, Lymer JM, Matta R, Clipperton-Allen AE, Kavaliers M, and Choleris E. Estrogen involvement in social behavior in rodents: Rapid and long-term actions. Horm Behav. 2015; 74: 53–76.

62. Hung YC, Wu YJ, Chien ME, Lin YT, Tsai CF, and Hsu KS. Loss of oxytocin receptors in hilar mossy cells impairs social discrimination. Neurobiol Dis. 2023; 187: 106311.

63. Scharfman HE, Hintz TM, Gomez J, Stormes KA, Barouk S, Malthankar-Phatak GH, McCloskey DP, Luine VN, and Maclusky NJ. Changes in hippocampal function of ovariectomized rats after sequential low doses of estradiol to simulate the preovulatory estrogen surge. Eur J Neurosci. 2007; 26: 2595–612.

64. Blaustein JD. An estrogen by any other name. Endocrinology. 2008; 149: 2697–8.

65. Oberlander JG and Woolley CS. 17β-estradiol acutely potentiates glutamatergic synaptic transmission in the hippocampus through distinct mechanisms in males and females. J Neurosci. 2016; 36: 2677–90.

66. Zadran S, Qin Q, Bi X, Zadran H, Kim Y, Foy MR, Thompson R, and Baudry M. 17β-estradiol increases neuronal excitability through MAP kinase-induced calpain activation. Proc Natl Acad Sci U S A. 2009; 106: 21936–41.

67. Rudick CN and Woolley CS. Estradiol induces a phasic Fos response in the hippocampal CA1 and CA3 regions of adult female rats. Hippocampus. 2000; 10: 274–83.

68. West AE, Pruunsild P, and Timmusk T. Neurotrophins: Transcription and translation. Handb Exp Pharmacol. 2014; 220: 67–100.

69. Bouchenak M and Lamri-Senhadji M. Nutritional quality of legumes, and their role in cardiometabolic risk prevention: A review. J Med Food. 2013; 16: 185–98.

70. Hloucalova P, Skladanka J, Horky P, Klejdus B, Pelikan J, and Knotova D. Determination of phytoestrogen content in fresh-cut legume forage. Animals (Basel). 2016; 6.

71. Hsiao K, Chapman P, Nilsen S, Eckman C, Harigaya Y, Younkin S, Yang F, and Cole G. Correlative memory deficits, aβ elevation, and amyloid plaques in transgenic mice. Science. 1996; 274: 99–102.

72. Smith MS, Freeman ME, and Neill JD. The control of progesterone secretion during the estrous cycle and early pseudopregnancy in the rat: Prolactin, gonadotropin and steroid levels associated with rescue of the corpus luteum of pseudopregnancy. Endocrinology. 1975; 96: 219–26.

73. Whitten WK. Occurrence of anoestrus in mice caged in groups. J Endocrinol. 1959; 18: 102–7.

74. Scharfman HE, Kim M, Hintz TM, and MacLusky NJ. Seizures and reproductive function: Insights from female rats with epilepsy. Ann Neurol. 2008; 64: 687–97.

75. Criscuolo C, Chartampila E, Ginsberg SD, and Scharfman HE. Stability of dentate gyrus granule cell mossy fiber BDNF protein expression with age and resistance of granule cells to Alzheimer’s disease neuropathology in a mouse model. eNeuro. 2023; 11: ENEURO.0192-23.2023.

76. Danielson NB, Turi GF, Ladow M, Chavlis S, Petrantonakis PC, Poirazi P, and Losonczy A. In vivo imaging of dentate gyrus mossy cells in behaving mice. Neuron. 2017; 93: 552–559 e4.

77. Leranth C, Szeidemann Z, Hsu M, and Buzsaki G. AMPA receptors in the rat and primate hippocampus: A possible absence of GluR2/3 subunits in most interneurons. Neuroscience. 1996; 70: 631–52.

78. Gonzalez-Gomez M and Meyer G. Dynamic expression of calretinin in embryonic and early fetal human cortex. Front Neuroanat. 2014; 8: 41.

79. Vieillard J, Franck MCM, Hartung S, Jakobsson JET, Ceder MM, Welsh RE, Lagerstrom MC, and Kullander K. Adult spinal Dmrt3 neurons receive direct somatosensory inputs from ipsi- and contralateral primary afferents and from brainstem motor nuclei. J Comp Neurol. 2023; 531: 5–24.

80. Botterill JJ, Gerencer KJ, Vinod KY, Alcantara-Gonzalez D, and Scharfman HE. Dorsal and ventral mossy cells differ in their axonal projections throughout the dentate gyrus of the mouse hippocampus. Hippocampus. 2021; 31: 522–539.

81. Alcantara-Gonzalez D, Kennedy M, Criscuolo C, Botterill J, and Scharfman HE. Increased excitability of dentate gyrus mossy cells occurs early in life in the Tg2576 model of Alzheimer’s disease. Alzheimers Res Ther. 2025; 17: 105.

82. Linse S, Voorhies M, Norstrom E, and Schultz DA. An EF-hand phage display study of calmodulin subdomain pairing. J Mol Biol. 2000; 296: 473–86.

83. Turner EC, Gabi M, Liao CC, and Kaas JH. The postnatal development of MT, V1, LGN, pulvinar and SC in prosimian galagos (otolemur garnettii). J Comp Neurol. 2020; 528: 3075–3094.

84. Zimmermann L and Schwaller B. Monoclonal antibodies recognizing epitopes of calretinins: Dependence on Ca2+-binding status and differences in antigen accessibility in colon cancer cells. Cell Calcium. 2002; 31: 13–25.

85. Huang Y, Zhang L, Song NN, Hu ZL, Chen JY, and Ding YQ. Distribution of satb1 in the central nervous system of adult mice. Neurosci Res. 2011; 71: 12–21.

86. Mundrucz L, Kecskes A, Henn-Mike N, Kobor P, Buzas P, Vennekens R, and Kecskes M. TRPM4 regulates hilar mossy cell loss in temporal lobe epilepsy. BMC Biol. 2023; 21: 96.

87. Herbison AE. Neurochemical identity of neurones expressing oestrogen and androgen receptors in sheep hypothalamus. J Reprod Fertil Suppl. 1995; 49: 271–83.

88. Price RH, Jr. and Handa RJ. Expression of estrogen receptor-β protein and mRNA in the cerebellum of the rat. Neurosci Lett. 2000; 288: 115–8.

89. Stenberg AE, Simonoska R, Stygar D, Sahlin L, and Hultcrantz M. Effect of estrogen and antiestrogens on the estrogen receptor content in the cochlea of ovariectomized rats. Hear Res. 2003; 182: 19–23.

90. Skinner DC and Dufourny L. Oestrogen receptor β-immunoreactive neurones in the ovine hypothalamus: Distribution and colocalisation with gonadotropin-releasing hormone. J Neuroendocrinol. 2005; 17: 29–39.

91. Zhang Y, Xiao X, Zhang XM, Zhao ZQ, and Zhang YQ. Estrogen facilitates spinal cord synaptic transmission via membrane-bound estrogen receptors: Implications for pain hypersensitivity. J Biol Chem. 2012; 287: 33268–81.

92. Meseke M, Neumuller F, Brunne B, Li X, Anstotz M, Pohlkamp T, Rogalla MM, Herz J, Rune GM, and Bender RA. Distal dendritic enrichment of HCN1 channels in hippocampal CA1 is promoted by estrogen, but does not require reelin. eNeuro. 2018; 5.

93. Wu Y, Feng D, Lin J, Qu Y, He S, Wang Y, Gao G, and Zhao T. Downregulation of g-protein-coupled receptor 30 in the hippocampus attenuates the neuroprotection of estrogen in the critical period hypothesis. Mol Med Rep. 2018; 17: 5716–5725.

94. Zhao J, Bian C, Liu M, Zhao Y, Sun T, Xing F, and Zhang J. Orchiectomy and letrozole differentially regulate synaptic plasticity and spatial memory in a manner that is mediated by SRC-1 in the hippocampus of male mice. J Steroid Biochem Mol Biol. 2018; 178: 354–368.

95. Broughton BR, Brait VH, Guida E, Lee S, Arumugam TV, Gardiner-Mann CV, Miller AA, Tang SC, Drummond GR, and Sobey CG. Stroke increases g protein-coupled estrogen receptor expression in the brain of male but not female mice. Neurosignals. 2013; 21: 229–39.

96. Amaral DG. A Golgi study of cell types in the hilar region of the hippocampus in the rat. J Comp Neurol. 1978; 182: 851–914.

97. Amaral DG, Scharfman HE, and Lavenex P. The dentate gyrus: Fundamental neuroanatomical organization (dentate gyrus for dummies). Prog Brain Res. 2007; 163: 3–22.

98. Chen ZK, Quintanilla L, Su Y, Sheehy RN, Simon JM, Luo YJ, Li YD, Chen Z, Asrican B, Tart DS, Farmer WT, Ming GL, Song H, and Song J. Septo-dentate gyrus cholinergic circuits modulate function and morphogenesis of adult neural stem cells through granule cell intermediaries. Proc Natl Acad Sci U S A. 2024; 121: e2405117121.

99. Llorente R, Marraudino M, Carrillo B, Bonaldo B, Simon-Areces J, Abellanas-Perez P, Rivero-Aguilar M, Fernandez-Garcia JM, Pinos H, Garcia-Segura LM, Collado P, and Grassi D. G protein-coupled estrogen receptor immunoreactivity fluctuates during the estrous cycle and show sex differences in the amygdala and dorsal hippocampus. Front Endocrinol (Lausanne). 2020; 11: 537.

100. Levin ER. Plasma membrane estrogen receptors. Trends Endocrinol Metab. 2009; 20: 477–82.

101. McEwen BS and Milner TA. Understanding the broad influence of sex hormones and sex differences in the brain. J Neurosci Res. 2017; 95: 24–39.

102. Pillerova M, Borbelyova V, Hodosy J, Riljak V, Renczes E, Frick KM, and Tothova L. On the role of sex steroids in biological functions by classical and non-classical pathways. An update. Front Neuroendocrinol. 2021; 62: 100926.

103. Plant TM, Zeleznik AJ, and Albertini DF, eds. Knobil and neill’s physiology of reproduction. 4th ed. 2014, Elsevier: New York.

104. Diniz DM, Calabrese F, Brivio P, Riva MA, Grandjean J, and Homberg JR. BDNF overexpression in the ventral hippocampus promotes antidepressant- and anxiolytic-like activity in serotonin transporter knockout rats. Int J Mol Sci. 2021; 22.

105. Gualtieri F, Bregere C, Laws GC, Armstrong EA, Wylie NJ, Moxham TT, Guzman R, Boswell T, and Smulders TV. Effects of environmental enrichment on doublecortin and BDNF expression along the dorso-ventral axis of the dentate gyrus. Front Neurosci. 2017; 11: 488.

106. Piatti VC and Schinder AF. Hippocampal mossy cells provide a fate switch for adult neural stem cells. Neuron. 2018; 99: 425–427.

107. Yeh CY, Asrican B, Moss J, Quintanilla LJ, He T, Mao X, Casse F, Gebara E, Bao H, Lu W, Toni N, and Song J. Mossy cells control adult neural stem cell quiescence and maintenance through a dynamic balance between direct and indirect pathways. Neuron. 2018; 99: 493–510 e4.

108. Scharfman HE and Bernstein HL. Potential implications of a monosynaptic pathway from mossy cells to adult-born granule cells of the dentate gyrus. Front Syst Neurosci. 2015; 9: 112.

109. Eide FF, Lowenstein DH, and Reichardt LF. Neurotrophins and their receptors--current concepts and implications for neurologic disease. Exp Neurol. 1993; 121: 200–14.

110. Davies AM. The neurotrophic hypothesis: Where does it stand? Philos Trans R Soc Lond B Biol Sci. 1996; 351: 389–94.

111. Lewin GR. Neurotrophins and the specification of neuronal phenotype. Philos Trans R Soc Lond B Biol Sci. 1996; 351: 405–11.

112. Wang CS, Kavalali ET, and Monteggia LM. BDNF signaling in context: From synaptic regulation to psychiatric disorders. Cell. 2022; 185: 62–76.

113. Blasco-Ibanez JM and Freund TF. Distribution, ultrastructure, and connectivity of calretinin-immunoreactive mossy cells of the mouse dentate gyrus. Hippocampus. 1997; 7: 307–20.

114. Fujise N, Liu Y, Hori N, and Kosaka T. Distribution of calretinin immunoreactivity in the mouse dentate gyrus: II. Mossy cells, with special reference to their dorsoventral difference in calretinin immunoreactivity. Neuroscience. 1998; 82: 181–200.

115. Liu Y, Fujise N, and Kosaka T. Distribution of calretinin immunoreactivity in the mouse dentate gyrus. I. General description. Exp Brain Res. 1996; 108: 389–403.

116. Jinno S, Ishizuka S, and Kosaka T. Ionic currents underlying rhythmic bursting of ventral mossy cells in the developing mouse dentate gyrus. Eur J Neurosci. 2003; 17: 1338–54.

117. Fredes F, Silva MA, Koppensteiner P, Kobayashi K, Joesch M, and Shigemoto R. Ventro-dorsal hippocampal pathway gates novelty-induced contextual memory formation. Curr Biol. 2021; 31: 25–38 e5.

118. Houser CR, Peng Z, Wei X, Huang CS, and Mody I. Mossy cells in the dorsal and ventral dentate gyrus differ in their patterns of axonal projections. J Neurosci. 2021; 41: 991–1004.

119. Huang CS and Houser CR. Alterations and imbalance of dorsal and ventral mossy cells in a mouse model of epilepsy. bioRxiv. 2024.

120. Barker JM and Galea LA. Sex and regional differences in estradiol content in the prefrontal cortex, amygdala and hippocampus of adult male and female rats. Gen Comp Endocrinol. 2009; 164: 77–84.

121. Frick KM, Kim J, and Koss WA. Estradiol and hippocampal memory in female and male rodents. Curr Opin Behav Sci. 2018; 23: 65–74.

122. Hokenson RE, Rodriguez-Acevedo KL, Chen Y, Short AK, Samrari SA, Devireddy B, Jensen BJ, Winter JJ, Gall CM, Soma KK, Heller EA, and Baram TZ. Hippocampal estrogen levels, receptor types, and epigenetics contribute to sex-specific memory vulnerabilities to concurrent acute stresses. Neuron. 2026; 114: 1454–1472 e8.

123. Brandt N, Vierk R, and Rune GM. Sexual dimorphism in estrogen-induced synaptogenesis in the adult hippocampus. Int J Dev Biol. 2013; 57: 351–6.

124. Kelly MJ, Loose MD, and Ronnekleiv OK. Estrogen suppresses μ-opioid-and GABAB-mediated hyperpolarization of hypothalamic arcuate neurons. J Neurosci. 1992; 12: 2745–50.

125. Bendis PC, Zimmerman S, Onisiforou A, Zanos P, and Georgiou P. The impact of estradiol on serotonin, glutamate, and dopamine systems. Front Neurosci. 2024; 18: 1348551.

126. Luine VN, Richards ST, Wu VY, and Beck KD. Estradiol enhances learning and memory in a spatial memory task and effects levels of monoaminergic neurotransmitters. Horm Behav. 1998; 34: 149–62.

127. Yu M, Yin N, Feng B, Gao P, Yu K, Liu H, Liu H, Li Y, Ginnard OZ, Conde KM, Wang M, Fang X, Tu L, Bean JC, Liu Q, Deng Y, Yang Y, Han J, Jossy SV, Burt ML, Wong HZ, Yang Y, Arenkiel BR, He Y, Guo S, Gourdy P, Arnal JF, Lenfant F, Wang Z, Wang C, He Y, and Xu Y. Identification of an ionic mechanism for ERα-mediated rapid excitation in neurons. Sci Adv. 2024; 10: eadp0696.

128. Jain A and Woolley CS. Mechanisms that underlie expression of estradiol-induced excitatory synaptic potentiation in the hippocampus differ between males and females. J Neurosci. 2023; 43: 1298–1309.

129. Kramar EA, Chen LY, Rex CS, Gall CM, and Lynch G. Estrogen’s place in the family of synaptic modulators. Mol Cell Pharmacol. 2009; 1: 258–262.

130. Fester L and Rune GM. Sex neurosteroids: Hormones made by the brain for the brain. Neurosci Lett. 2021; 753: 135849.

131. Win R, Minto W, Mah IK, and Boyd K. Integration of chromogenic rnascope in situ hybridization for target validation in drug discovery. Toxicol Pathol. 2025; 53: 21–30.

132. Scharfman HE and Maclusky NJ. Similarities between actions of estrogen and BDNF in the hippocampus: Coincidence or clue? Trends Neurosci. 2005; 28: 79–85.

133. Cooke BM and Woolley CS. Gonadal hormone modulation of dendrites in the mammalian cns. J Neurobiol. 2005; 64: 34–46.

134. McEwen B. Estrogen actions throughout the brain. Recent Prog Horm Res. 2002; 57: 357–84.

135. Hoffman GE, Moore N, Fiskum G, and Murphy AZ. Ovarian steroid modulation of seizure severity and hippocampal cell death after kainic acid treatment. Exp Neurol. 2003; 182: 124–34.

136. Veliskova J, Velisek L, Galanopoulou AS, and Sperber EF. Neuroprotective effects of estrogens on hippocampal cells in adult female rats after status epilepticus. Epilepsia. 2000; 41 Suppl 6: S30–5.

